# Mice are not automatons; subjective experience in premotor circuits guides behavior

**DOI:** 10.1101/2021.06.23.449617

**Authors:** Drew C. Schreiner, Christian Cazares, Rafael Renteria, Christina M. Gremel

## Abstract

Subjective experience is a powerful driver of decision-making and continuously accrues. However, most neurobiological studies constrain analyses to task-related variables and ignore how continuously and individually experienced internal, temporal, and contextual factors influence adaptive behavior during decision-making and the associated neural mechanisms. We show mice rely on learned information about recent and longer-term subjective experience of variables above and beyond prior actions and reward, including checking behavior and the passage of time, to guide self-initiated, self-paced, and self-generated actions. These experiential variables were represented in secondary motor cortex (M2) activity and its projections into dorsal medial striatum (DMS). M2 integrated this information to bias strategy-level decision-making, and DMS projections used specific aspects of this recent experience to plan upcoming actions. This suggests diverse aspects of experience drive decision-making and its neural representation, and shows premotor corticostriatal circuits are crucial for using selective aspects of experiential information to guide adaptive behavior.

## Introduction

Most neurobiological investigations into decision-making seek to use well-constrained tasks to isolate specific components of decision-making and illuminate the corresponding neural mechanisms. These investigations often institute a trial structure, limit choice and movement, and elicit behavior via cues, with the latter leading to a historical focus on elicited stimulus-response characterization of involved mechanisms (Juavinett et al., 2018). Thus, interpretation of the associated mechanisms and models are made within this well-constrained vacuum. There is growing concern that such an approach negates the very individualistic, experiential, and continuous nature of decision-making (Balleine, 2019; Gomez-Marin et al., 2014; Krakauer et al., 2017; Schreiner et al., 2021; Yoo et al., 2021). Presumably, the continuous experiential information accrued by the self is reflected in and used by the brain to execute adaptive behavior to support ongoing decision-making. Yet such information is often treated as task-irrelevant and ignored or factored out (Roy et al., 2021), to an at best incomplete, or at worst inaccurate picture of involved neural mechanisms. Indeed, the seemingly widespread distribution of similar decision-making information across the brain (Allen et al., 2017; Steinmetz et al., 2019) may in part be due to a lack of accounting for the unique constellation of internal, experiential, temporal, and contextual information encountered by an individual that drives decision-making, referred to here as “subjective experience”.

Evidence from ethological approaches shows that diverse types of subjective experience contribute to decision-making. When dropping shelled prey to break them, a crow will integrate the type of prey, the number of times the item has been dropped, the hardness of the surface, and the amount of kleptoparasitism to determine how high to drop the item (Cristol and Switzer, 1999). Behavior need not be complex; even innate behaviors are modified by experience (Remedios et al., 2017) and simple experience-based strategies such as win-stay and lose-shift persist after performance (a proxy for learning) plateaus. Experiential-based emergence of control does not happen within a vacuum; an individual’s interactions with their environment drive adaptive behavior (Balleine, 2019; Costa, 2011) with contributions from temporal (Ariely and Zakay, 2001), historical and contextual (Bouton and Balleine, 2019), and internal state factors (Balleine and Dickinson, 1998; Berridge et al., 2008). Indeed, there is an interplay between exploration and the accrual of experiential information, with exploration uncovering contingency and consequence information that in turn can be used to bias towards further exploration or experience-guided decision-making. However, in many investigations, experience is either not needed or even actively detrimental to performance (e.g., in perceptual decision-making tasks, subjects should ideally attend only to the current stimulus). That subjective experience appears to contribute even where it is “unnecessary” argues that it is a powerful driver of adaptive behavior (Lak et al., 2020). Further, many investigations constrain behavior by using trial-based, binary decision tasks unlikely to generalize fully to more ethological, self-generated, and continuously evolving types of decision-making (Yoo et al., 2021).

As subjective experience can play a large role in psychiatric disease (e.g., the temporal pattern of drug use is decisive in substance use disorders (Allain et al., 2015)), there is a need to account for its influence on the behavioral and neural mechanisms of decision-making. One neural circuit that is disrupted in disease and presents as a candidate for the integration of such experiential information is secondary motor cortex (M2) (Ebbesen et al., 2018). On one hand, M2’s sensory, motor, and premotor characteristics have implicated a role in using experience to guide decision-making (Erlich et al., 2011; Murakami et al., 2014; Pinto et al., 2019; Siniscalchi et al., 2016). On the other hand, several studies have found that M2 appears to be involved in implementing stochastic or exploratory decisions (Murakami et al., 2017; Pisupati et al., 2021; Schreiner and Gremel, 2018; Tervo et al., 2014). However, animals may decide to explore *based upon their experience*; for instance making more exploratory decisions when uncertainty is high (Dhawale et al., 2019). Thus, attribution of M2 function to seemingly disparate processes may reflect the lack of accounting for or limiting the contribution of experiential information. Instead, we hypothesize that M2 represents and integrates experiential information to guide experience or exploration-based decision-making when use of such information is advantageous. This strategy-level control over action selection may be exerted through M2 projections into dorsal medial striatum (DMS) (Delevich et al., 2020; Hintiryan et al., 2016). Indeed, recently M2-DMS projections have been implicated in repetitive actions in a mouse model of Obsessive Compulsive Disorder (Corbit et al., 2019), suggesting that M2-DMS dysfunction may contribute to disease states. In order to capture the contribution of experiential information to decision-making and its potential neural implementation, we need to investigate within a framework where the use of experience is *essential* and not merely incidental to decision-making. Here, we utilized an unstructured free operant foraging task with continuous variables in mice where experience is crucial for efficient performance. We find aspects of experiential information normally considered task-irrelevant play large roles in supporting adaptive behavior, often more than that played by reward itself. We then show M2 circuits and their output to dorsal medial striatum (DMS) are crucial for use of this subjective experience to drive adaptive behavior.

## Results

### Mice learned an unstructured, self-generated, self-paced lever press hold down task

We adapted an instrumental task (Fan et al., 2012; Platt et al., 1973; Yin, 2009) where mice (n = 12 C57BL/6J) were trained to press and hold down a lever for at least a minimum duration to earn a food reward, with reward delivered at lever press release/offset (Figure 1A). There were no external cues signaling reward availability or duration, nor any trial structure (lever was always available). Thus lever presses were self-initiated, self-paced, and self-terminated and mice had to explore the contingency duration to determine the rule, a process termed action differentiation (Skinner, 1938).

**Figure 1.**
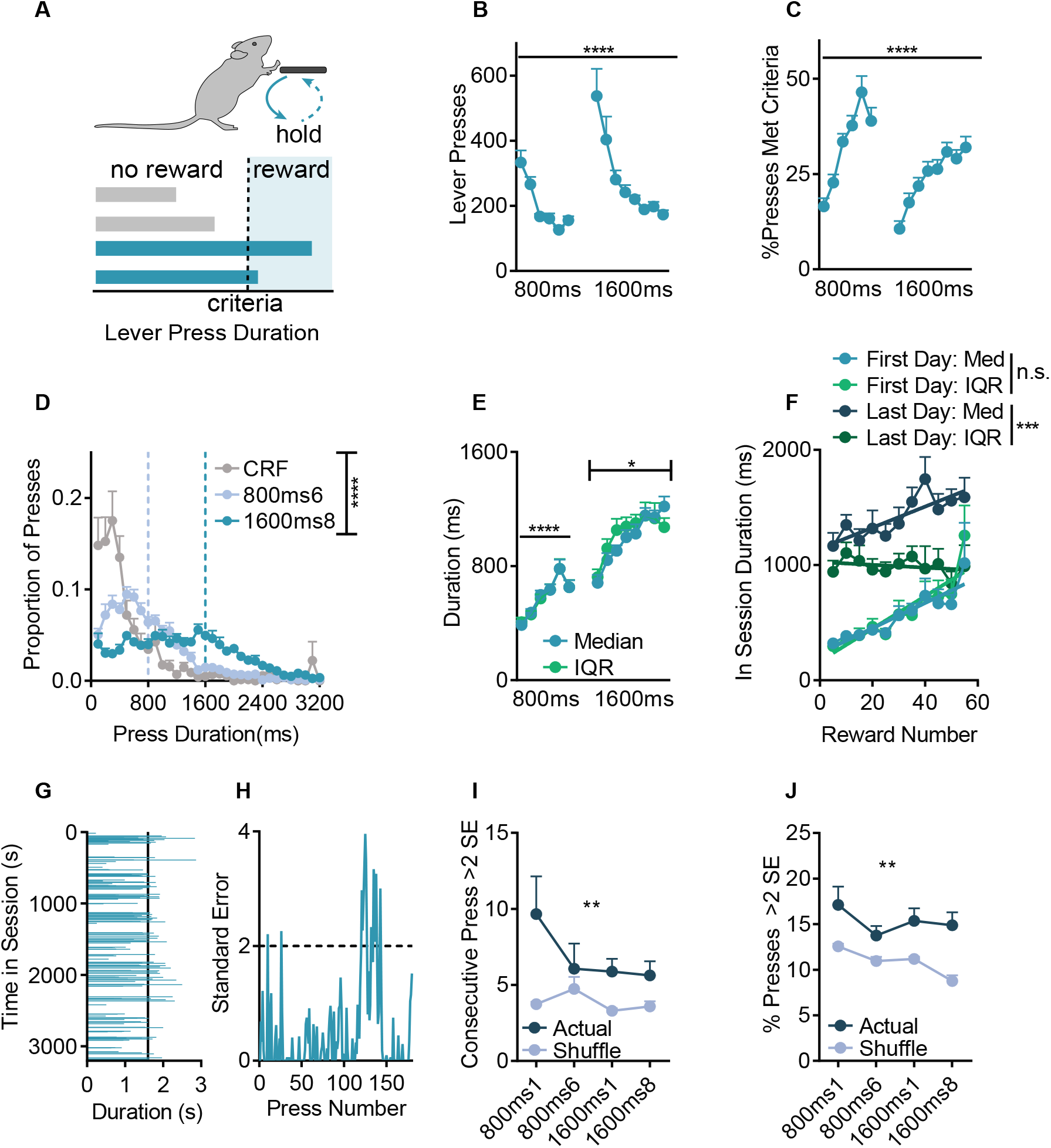
Mice learned an unstructured, self-generated, self-paced lever press hold down task. Behavioral schematic; mice learn to press and hold down a lever for at least a minimum duration to earn food reward. (B) Total Lever Presses across training days. (C) %Presses that met criteria across training. (D) Histogram of lever press durations (100ms bins) on the final pretraining day (CRF = Continuous Ratio of Reinforcement), and the final 800ms and 1600ms days. Dashed lines indicate criterion. (E) Median and Interquartile Range (IQR) of lever press durations across training days. (F) Duration median (Med) and IQR within a session, grouped by cumulative number of rewards. (G) Sample behavior of one trained mouse showing press durations in order of occurrence. (H) Upper cumulative sum from the same mouse/session. (I) Number of consecutive presses and (J) Overall % of presses that were >2 Standard Errors (SE) above the mean in the upper cumulative sum. 800ms and 1600ms refer to days where criterion was >800ms or >1600ms. **** p < 0.0001, *** p < 0.001, ** p < 0.01, * p < .05. Points represent mean+SEM across mice, unless noted otherwise.

We first examined macroscopic aspects of lever pressing. Mice were initially trained with an >800ms criterion before being shifted to a >1600ms criterion. Mice readily learned that press duration was the operant and quickly reduced the number of Total Lever Presses (Figure 1B; 1-way ANOVA F_2.9, 31.9_ = 12.0, p < 0.0001), while they increased the percentage of presses that met the minimum duration criterion (Figure 1C; referred to as %Presses Met Criteria, 1-way ANOVA F_4.22, 46.5_ = 17.2, p < 0.0001), and showed little evidence of stereotypies in their lever pressing (see Note S1). Mice were sensitive to the minimum duration rule and shifted the distribution of press durations from a pretraining session with no duration requirement, to the final day of >800ms training, and further still to the final day of >1600ms training (Figure 1D; 2-way RM ANOVA, main effect of Duration Bin F_31,1056_ = 34.1, p < 0.0001, and an interaction (Duration Bin/Criterion) F_62,1056_ = 10.5, p < 0.0001). To examine whether actions were controlled by their expected consequence and operationally goal-directed or were instead habitual (Adams and Dickinson, 1981; Dickinson, 1985), we performed outcome devaluation testing (Figure S1A). Mice reduced their Total Lever Presses on Devalued days relative to Valued days (Figure S1B), consistent with using expected outcome value to guide decisions as seen in goal-directed control (Adams and Dickinson, 1981). Although Total Lever Presses decreased, the %Presses Met Criteria *increased* following devaluation (Figure S1C) with a small rightward shift in the distribution of press durations (Figure S1D), suggesting action selection and execution may be differentially controlled by outcome value.

It is clear that mice can use contingency and consequence information to perform this task, but it is unclear *how* they are doing so. One possibility is that executed lever press durations are independent, with mice timing each press. If so, we hypothesized that mice may exhibit the scalar property of timing; as lever press durations increase, so too does variability (Gibbon et al., 1984; Yin, 2009). We calculated the median and interquartile range (IQR) of each animal’s lever press durations across training (Figure 1E) and found concomitant increases in both the median and the IQR across training days during initial short criterion training (2-way RM ANOVA, main effect only of Day (F_5,55_ = 19.5, p < 0.0001). However, when the duration criterion increased and training continued, the pattern of change in lever press IQR departed from the pattern of change in lever press median duration (2-way RM ANOVA, main effect of Day F_7,77_ = 14.0, p < 0.0001, and an interaction (Median/IQR x Day) F_7,77_ = 2.44, p = 0.026), suggesting reduced reliance on timing information. Reminiscent of skill learning, mice also showed within session increases in median durations during both the first and *last* day of training (Figure 1F). Linear regressions showed a significantly non-zero slope on both the first >800ms day (F_1,110_ = 28.9, p < 0.0001, R^2^ = 0.21) as well as the final >1600ms day (F_1,115_ = 12.6, p = 0.0006, R^2^ = 0.099). However, although within session increases in IQR were present on the first day of training (F_1,110_ = 48.5, p < 0.0001, R^2^ = 0.306), by the final day of training IQR no longer increased within a session (F_1,115_ = 0.28, p = 0.59, R^2^ = 0.002). Furthermore, while the slopes of the within session median and IQR were not different on the first day of training (p = 0.27), they *were* different by the final day (F_1,230_ = 9.1, p = 0.003), in violation of the scalar property of timing. This suggests mice used additional information other than solely timing behavior to control lever pressing.

What is this non-timing information? One possibility is that mice relied on recent experience to guide their decision-making. In Figure 1G, we plot press durations across one session for one well-trained mouse. Immediately clear is the rich behavior, both in terms of when presses occurred, and in their duration. However, there also appeared to be distinct periods of reduced variability. A cumulative sum (upper bound) analysis (Figure 1H) uncovered prolonged periods of time when mice emitted press durations >2 standard errors (SE) above the mean, (Figures 1I-J). This was not due to random chance, or the consequence of very long press durations inflating the cumulative sum (2-way RM ANOVA; percentage of >2SE presses: main effect of Day F_3,33_ = 3.98, p = 0.016, main effect of Actual vs. Shuffled F_1,11_ = 17.1, p = 0.0017. Consecutive >2SE presses: main effect only of Actual vs. Shuffled F_1,11_ = 14.0, p = 0.0032).

### Subjective experience contributes to internally-generated decision-making

The relative similarity among serial lever presses suggests that recent experience may contribute to adaptive behavior in support of self-generated decision-making. We modeled the effect of recent press history on performance by creating a simple linear mixed effect model (LME) that sought to predict current press duration (n) given recently executed durations (n-back). We included random effects of both training day and mouse to account for the repeated structure of our data. We also included several control variables and compared the actual coefficients to those obtained from order shuffled data using permutation tests (STAR methods, Table S1). We found a consistent significant linear relationship between current press n duration and the durations of n - 1 through n - 6 presses, with the magnitude of this relationship decaying across n-back presses (Figure 2B). This suggests that recent subjective experience contributes to continuous decision-making.

**Figure 2.**
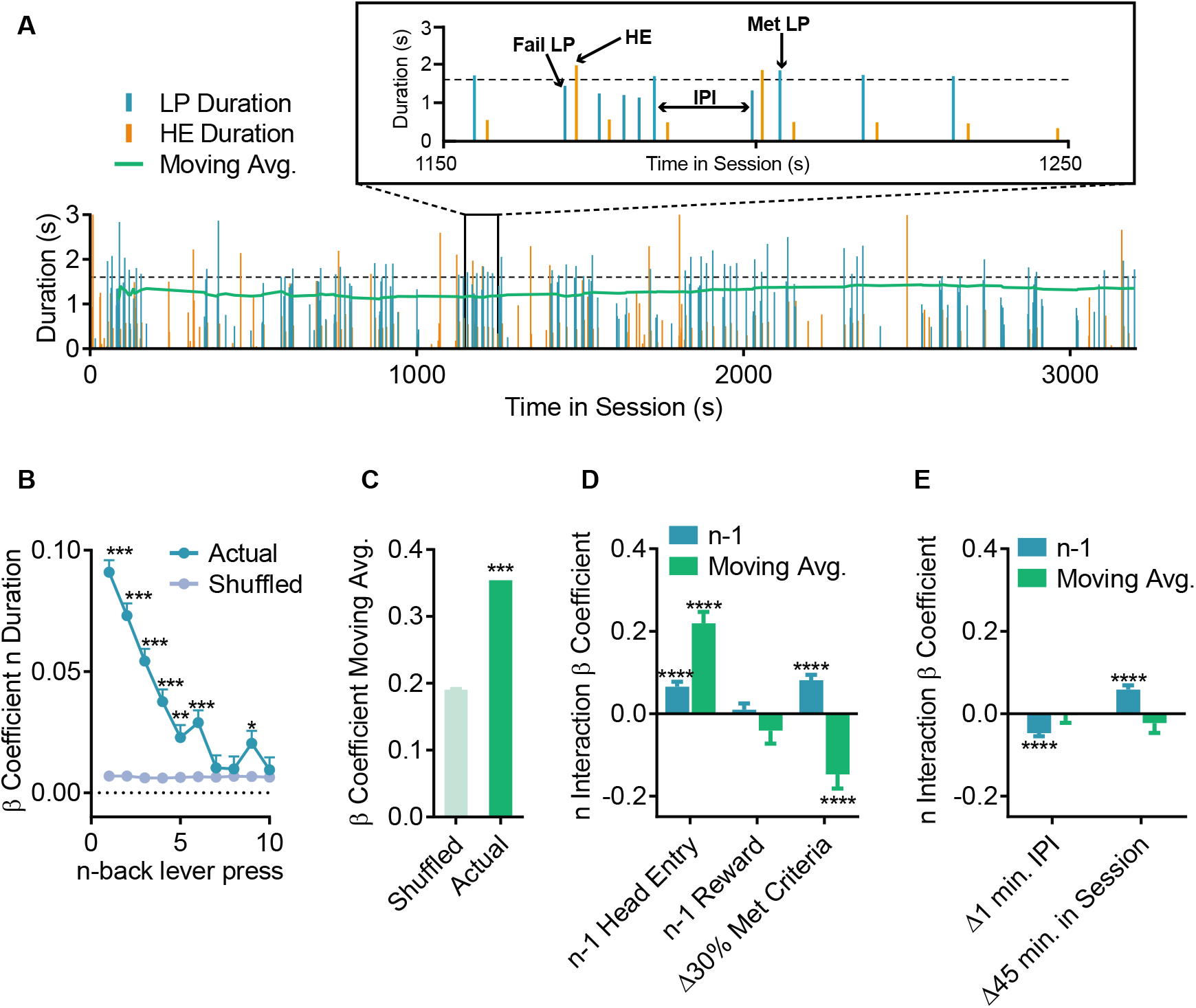
Subjective experience contributes to internally-generated decision-making. (A) Sample data from one mouse (as in Figures 1G-H) showing the diversity of experiential information available. Top shows a zoomed in subset. Dashed line indicates 1600ms criterion. β coefficients of LME model relating current lever press duration (n) to preceding press durations (n - x) for Actual and order Shuffled data. (C) Moving average β coefficient for Actual and Shuffled data. (D-E) β coefficients for the interaction between experiential variables and recent (n - 1 duration) or long-term (moving average) experience. For display purposes we transformed continuous variables to show relevant changes, e.g. time in session, which is in units of ms, was transformed to 45 min or half the duration of a session. LP = Lever Press, HE = Headentry into food magazine, IPI = Inter Press Interval. Δ = Change. * Markers in D-E indicate significant F-tests on model terms. **** p < 0.0001, *** p < 0.001, ** p < 0.01, * p < .05. Data points are mean+SEM. Shuffled data are mean+SEM of 1000 order shuffled β coefficients. See also Figure S2 and Tables S1-S3.

However, recent lever presses are not the only experiential information available (Figure 2A). The unstructured nature of this self-generated task allows us to capture aspects of decision-making that occur across a continuous space beyond just the press duration itself. We created more complex LMEs, first building a “full” model that included n - 1 through n - 6 durations, as well as main effect and interaction terms for other n-back variables, such as the inter-press interval between press n and press n - 1 (see Table S2 for terms). We performed backwards selection on this full model using Bayesian Information Criterion (BIC), leaving us with the model in Table S3. Follow-up permutation tests found that all the variables identified by BIC selection also significantly differed from order shuffled data (shuffled within a single mouse/session), suggesting that it is indeed the experienced order of these variables that affects subsequent press duration and not correlation across mice or across days. Importantly, when we built multiple linear regressions with the same experiential predictors, but using only individual mouse/session data, we found a strong correlation between model R^2^ and task performance (repeated measures correlation, R_m_ = 0.56, DF =153, p < 0.0001, slope = 0.38; Figure S2A). Thus, use of experience (as these models use only experience to predict duration) correlated with performance. Additionally, in the combined (all mice and training days) LMEs, there was a significant positive interaction between performance and the use of recent experience (serially adjacent presses were more similar in mice with higher performance, Figure 2D shows that an increase in %Met Criteria of 30% would roughly double the n/n - 1 relationship).

Of note, our goal with this model was not to make the most accurate predictions (though it predicted 24.1% of all lever press durations within a 95% CI, and accurately predicted whether a press did or did not meet criteria 73.8% of the time). Instead, we sought to ascertain 1) which experiential variables contributed, 2) if these variables interacted with previously made lever press durations, and 3) whether recent (i.e., n - 1) versus longer-term (i.e., a moving average of durations from n - 7 through n - 60) lever press duration experience differentially contributed (Iigaya et al., 2018). Beginning with the latter, we found that this long-term duration moving average coefficient significantly differed from order shuffled data (Figure 2C; permutation test p < 0.001). Thus, both recent and longer-term duration experience contributed to emitted durations.

### Observational, but not reward information contributes to decision-making

Experience can also be driven by information feedback processes. Here, headentry into the food magazine (HE) as a checking behavior provided information as to whether a lever press was or was not successful. HE behavior increased the relationship both between press n and n - 1, and between press n and the moving average (Figure 2D). The magnitude of this increase was quite large: lever presses within a lever press/HE/lever press sequence were effectively twice as related to one another relative to those in a lever press/lever press sequence. Thus checking behavior was a source of experiential information and influenced the subsequent executed behavior.

Success feedback may also be signaled by reward, and reward feedback is a crucial aspect of many decision-making and learning theories (e.g. Sutton and Barto, 1998; Rescorla and Wagner, 1972). Reward can modulate win-stay and lose-shift strategies even in well-learned tasks (Busse et al., 2011; Lak et al., 2020). However, it is less clear how reward might modulate decision-making in a more unconstrained task where one dictates their own behavioral opportunities. In regard to the present task, if mice “won and stayed”, we should expect that earning a reward on press n - 1 would cause mice to make a similar duration press afterwards.

We found no evidence of simple win-stay/lose-switch behavior, either in the use of recent (n - 1), or long-term (moving average) experience (Figure 2D). Put another way, earning a reward did not increase the similarity between press n and press n - 1, nor did failing to earn a reward lead to drastic shifts in behavior. We reasoned that perhaps the lack of a win-stay effect may have been due to reward being deterministic, and thus, only errors of execution could occur (McDougle et al., 2019). Therefore, we imposed a probabilistic reward schedule in a separate cohort (25%, 50%, or 75% rewarded, n = 5 mice per group) following training. Here, the %Met Criteria increased (Figure S2B). A Met Criteria press - whether or not it was rewarded - led to an increased relationship between press n and n - 1 (significant positive interaction, Figure S2C). The magnitude of this effect was *larger* when the Met press was unrewarded, and this “win-stay” effect was more pronounced in the groups where a Met press was least likely to produce a reward. Thus, mice made a press that was more similar to the one that preceded it after a Met press, especially if that press happened to be unrewarded due to chance. This provides evidence mice used an internal representation of press duration to guide behavior and relied less on the presence of reward.

### Time contributes to and modifies use of subjective experience in decision-making

The above findings challenge the assumption that decision-making is solely determined by the serial order of actions and their outcome, as is often presumed in trial-based experimental designs. That sources of this crucial experiential information, such as checking behaviors, accrue across a continuous temporal space raises the question of how the passage of time itself may influence decision-making. We find the relationship between two adjacent presses (presses n and n - 1) decreased as the inter-press-interval (IPI) increases (Figure 2E). To give an example of the magnitude, the model predicts that the relationship between n and n - 1 would be approximately 0 if they are separated by 120s. This raises the hypothesis that animals may rely more on the long-term moving average to guide their behavior following long IPIs (Iigaya et al., 2018). Indeed, the use of long-term experience was unaffected by the IPI. Further, we found that n and n - 1 became more similar towards the end of a session, and again, there was no relationship between time in session and the moving average (Figure 2E). Collectively, these results suggest that the passage of time is a crucial aspect to modifying recent experience, with less effect on the contribution of long-term learned contingencies.

### M2 represents prior experience and may guide exploration

M2 has been reported to be involved in both exploration and experience-based decision-making, and this apparent discrepancy may be due to neglecting the contribution of some of these checking, contextual, and temporal variables. Therefore, we performed pretraining lesions of M2 using ibotenic acid (Figure 3A; Lesion n = 10) or vehicle (Sham n = 8). In line with prior reports (Yin, 2009) we found no differences between Sham and Lesion mice in coarse behavioral measurements such as Total Lever Presses (Figure 3B), %Presses Met Criteria (Figure 3C), or Press Durations (Figure 3D). However, M2 lesioned mice executed lever press durations that were more similar to their prior action (Figure 3E). This was evidenced by a specific increase in the magnitude of the n - 1 β coefficient compared to Sham mice (2-way ANOVA (n-back x Sham/Lesion) main effect of n-back (F_9,572740_ = 20.2, p < 0.0001) and Sham/Lesion (F_1,572740_ = 14.1, p = 0.0002) and significant interaction (F_9,572740_ = 6.33, p < 0.0001). Post-hoc testing revealed a significant group difference only at n - 1 (t_572740_ = 6.87, p < 0.0001), with no differences at further n-back presses, nor in the moving average term (Figure 3F). Using the complex LME model, M2 lesions disrupted all n - 1 duration interactions, including Reward, Checking, IPI, and Time in Session (Figure 3G, see also Table S4). Lesions did not affect moving average interactions. Collectively, this suggests M2 lesioned mice were relatively inflexible, akin to previous studies where M2 lesions biased use of habitual or model-free processes (Gremel and Costa, 2013), and were left to rely on the just-made action without integration of broad experiential information.

**Figure 3.**
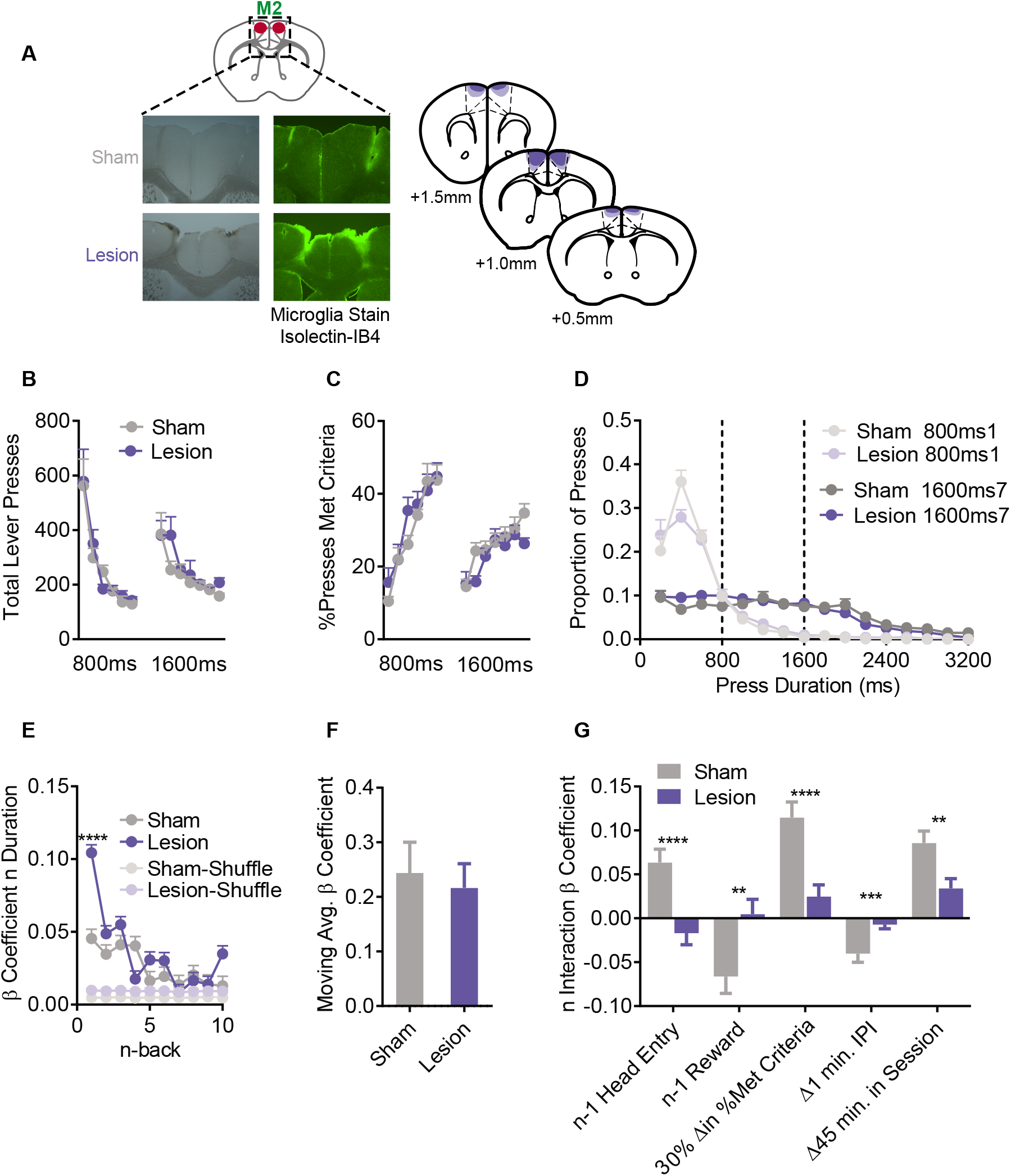
Pretraining lesions of M2 impair integration of experiential information. (A) (top left) Schematic, (bottom left) sample histology, and (right) average/maximal (dark/light shading) spread of excitotoxic lesions of M2, slice coordinates relative to Bregma. (B) Total Lever Presses across training days. (C) %Presses that met criteria across training days. (D) Histogram of lever press durations on the first and last days of the 800ms and 1600ms criteria (200ms bins). (E) β coefficient from LME models predicting n duration from n-back durations for Actual and order Shuffled data. (F) β coefficient for the moving average term. (G) β coefficients for the interaction terms from the complex LME model. For display purposes we transformed the continuous variables to show relevant changes. B-D are mean+SEM across mice. Shuffled data are the mean+SEM of 1000 order shuffled β coefficients. **** p < 0.0001, *** p < 0.001, ** p < 0.01. See also Table S4.

How may subjective experience influence representation of decision-making in M2 circuits? We utilized *in vivo* fiber photometry (Figure 4A; coordinates from Bregma: AP +1.0mm, L ±0.5mm and V −1.2mm, −1.4mm from the skull), and measured population Ca^2+^ activity from M2 excitatory neurons (n = 8 mice). Aligning baseline z-scored activity to lever press onset (Figure 4B), we observed preceding ramping activity as has been previously reported (Murakami et al., 2014). However, this M2 activity ramping did not differ based on whether that press would go on to exceed the criteria duration (Met) or not (Fail) (permutation testing that required 4 adjacent samples to pass the threshold for significance, see STAR methods and Jean-Richard-dit-Bressel et al., 2020). M2 activity during the lever press was modulated by whether that lever press would or would not meet the criteria (Figure 4C). This difference persisted following lever release, where there was an abrupt increase in Ca^2+^ activity just after the offset of Met presses, - i.e. reward delivery - as well as a subsequent sustained decrease in activity (Figure 4D). Thus M2 activity is modulated during lever pressing with ongoing modulation reflecting future success.

**Figure 4.**
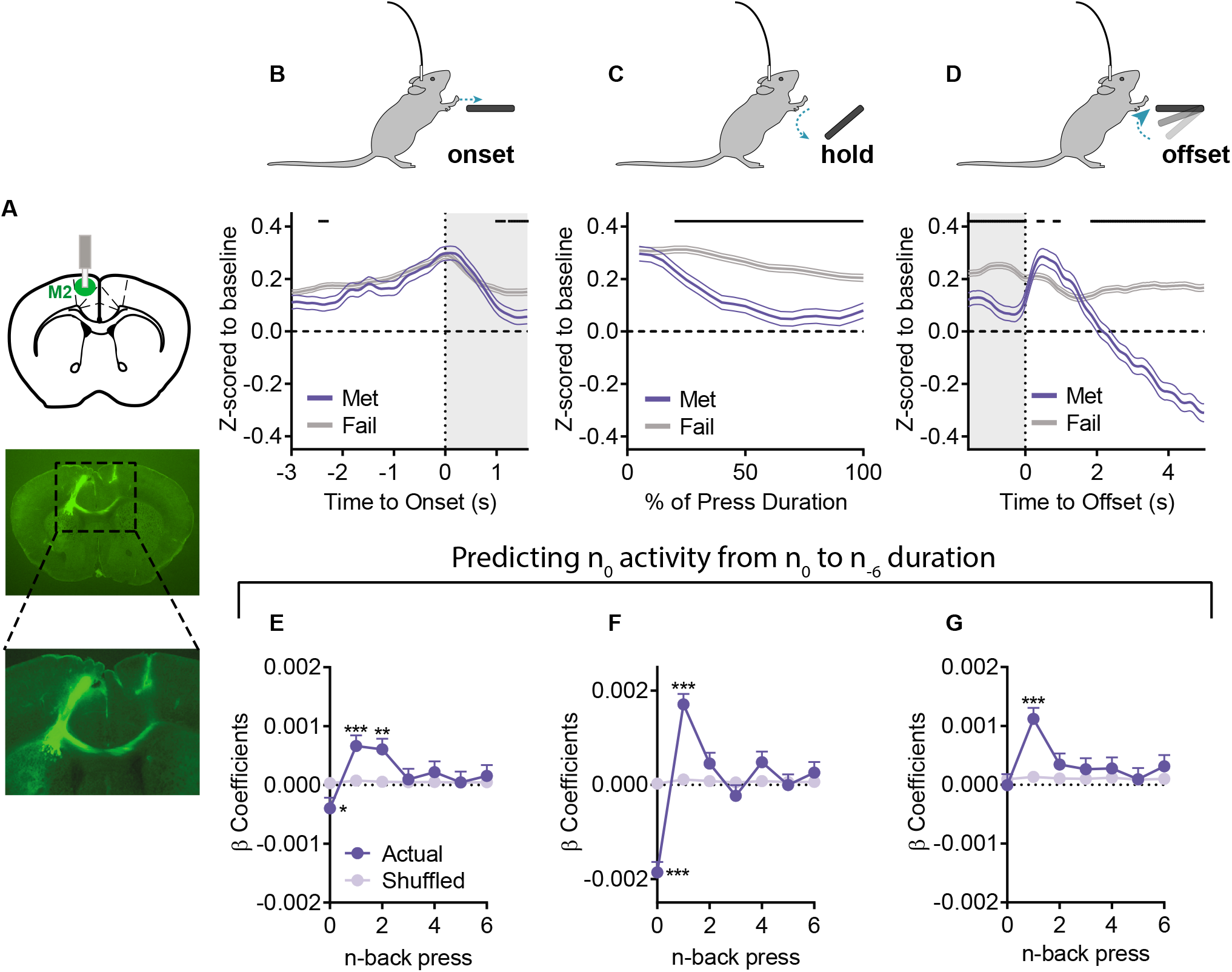
M2 Ca^2+^ activity represents prior experience. (A) (top) Schematic and (bottom) example histology of M2 *in vivo* Ca^2+^ fiber photometry recordings. (B-D) Ca^2+^ activity z-scored relative to a baseline period and aligned to (B) press onset, (C) the hold down period itself (presented as the relative % of a press’s duration), and (D) the offset of a lever press. (E-G) β coefficients from LME models relating activity to current and prior durations for Actual and order Shuffled data (E) before press onset, (F) during the press, and (G) after press offset. Met = Presses that met criterion, Fail = Presses that did not meet criterion. Grey shading in B, D indicates 1.6s. Black lines in B-D indicate significant differences between Met/Fail via permutation testing. Shuffled data are the mean+SEM of 1000 order shuffled β coefficients. *** p < 0.001, ** p < 0.01, * p < 0.05. See also Table S5.

To determine if M2 activity related to ongoing and prior behavior, we created LME models to predict Ca^2+^ activity during epochs of the current lever press given both the ongoing action (press n duration) and prior behavior (the duration of press n - 1 to n - 6). We included prior activity as a covariate to control for autocorrelation in Ca^2+^ activity and compared β coefficients to 1000 order shuffled datasets. Before the onset of press n, there was a significant positive relationship between M2 activity and the just prior press durations (Figure 4E; n - 1, p < 0.001; n - 2, p = 0.001) and a small negative relationship between activity and the upcoming duration (press n, p = 0.048). This representation of both current and prior lever press duration in M2 Ca^2+^ activity continued during the press itself (Figure 4F; press n, p < 0.001; n - 1: p < 0.001). At press offset, there was no relationship with the just completed press (n), but there was a positive relationship with the previous lever press duration (Figure 4G; n - 1, p < 0.001). We used complex LME models to investigate representation of other types of experiential information. M2 activity reflected Checking, IPI, and Time in Session, and these terms also interacted with the contribution of prior durations (Table S5). In particular, at each lever press epoch we observed a significant, positive interaction between n - 1 duration and HE checking in between press n and n - 1, suggesting that checking increased the relationship between lever press duration and M2 activity. These complex LMEs were also better at predicting M2 Ca^2+^ activity relative to the simple LMEs that only included durations (difference in simple/complex prediction %: Before Press: +13.7%, During Press: +9.2%, After Press: +16.4%), showing that these often neglected variables are powerful drivers of M2 activity. Thus, we see representation of diverse aspects of subjective experience, aspects whose contributions are lost when M2 is lesioned. This suggests M2 circuits are recruited when a broad array of experiential information is used to guide behavior (as is often seen in exploration), but not when behavior can be accomplished using less flexible (i.e. habitual/model-free) processes.

### M2-DMS projections use recent experience to plan upcoming actions

M2 sends dense projections into dorsal medial striatum (M2-DMS) regions (Delevich et al., 2020; Hintiryan et al., 2016) that contribute to action selection (Klaus et al., 2019), but it is unclear what information is conveyed. We performed *in vivo* fiber photometry of virally targeted M2-DMS activity and examined representation of experiential information within this population (n = 7, Figure 5A). We again observed a ramping in M2-DMS Ca^2+^ activity prior to lever press onset. This activity reflected future success, with larger increases in activity for presses that *would* meet criteria (Figure 5B). This relationship was also present during the press itself (Figure 5C), and upon lever release (Figure 5D), raising the hypothesis that M2-DMS projections may carry information specifying and/or planning actions based on prior experience.

**Figure 5.**
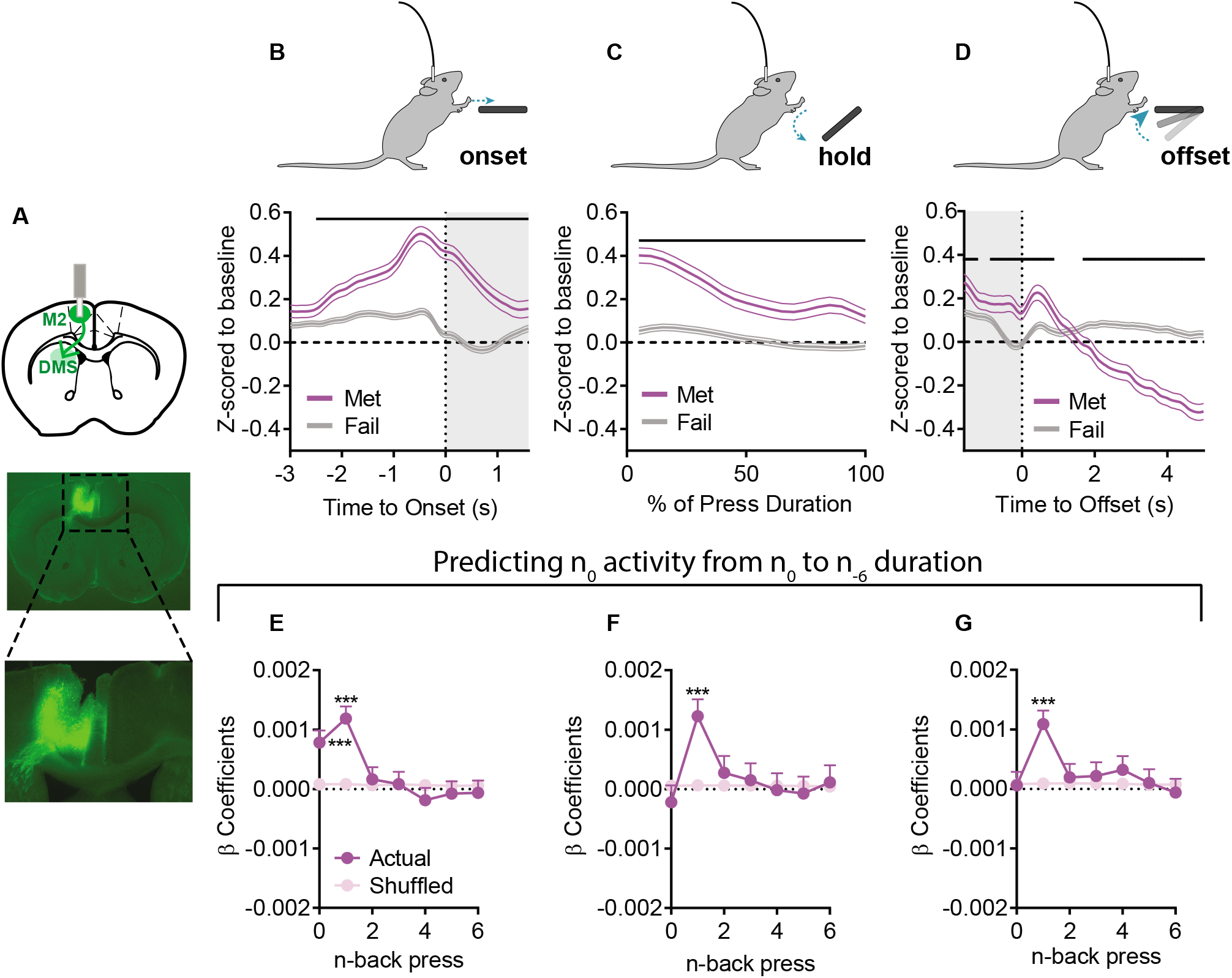
M2-DMS Ca^2+^ activity encodes preceding and upcoming actions. (A) (top) Schematic and (bottom) example histology of projection specific M2-DMS Ca^2+^ fiber photometry. (B-D) Ca^2+^ activity z-scored relative to baseline and aligned to (B) press onset, (C) the duration of the press, and (D) press offset. (E-G) β coefficients from LME models relating activity to current and prior durations for Actual and Shuffled data (E) before press onset, (F) during the press, and (G) after press offset. Met = Presses that met criteria. Fail = Presses that did not meet criterion. Grey shading in B, D indicates 1.6s. Black lines in B-D indicate significant differences between Met/Fail via permutation testing. Shuffled data are the mean+SEM of 1000 order shuffled β coefficients. *** p < 0.001. See also Table S6.

Indeed, LMEs using durations to predict M2-DMS activity before press onset showed both prior (Figure 5E; n - 1, p < 0.001) and upcoming (n, p < 0.001) durations were positively related to M2-DMS Ca^2+^ activity. M2-DMS activity during the press did not relate to the current duration, but was positively related to the prior duration (Figure 5F; n - 1, p < 0.001), and likewise at press offset there was a positive relationship only with the n - 1 press duration (Figure 5G; p < 0.001). Furthermore, complex LMEs revealed significant influences of Checking, prior Reward, IPI, and Time in Session on M2-DMS activity (Table S6). As in M2, at every time point there was an interaction between checking and n - 1 duration on M2-DMS activity. Further, the complex LMEs predicted more of the data relative to the simple models (difference in simple/complex prediction %: Before Press: +13.2%, During Press: +9.2%, After Press: +15.2%). Thus M2-DMS activity appears to reflect the use of recent experiences to plan upcoming actions.

To test whether M2-DMS activity functionally contributed to planning actions based on recent experience, we used a Cre-dependent caspase strategy to selectively lesion M2-DMS projection neurons prior to training (Figure 6A, n = 8 Lesion, n = 8 Sham). Again, we observed no effect on coarse measures of behavior including Total Lever Presses, %Presses Met Criteria, and Press Durations (Figures 6B-D). Simple LME modeling showed M2-DMS lesions reduced the relationship between press n and press n - 1 (Figure 6E; 2-way ANOVA (n-back x Sham/Lesion) main effect of n-back (F_9,467760_= 14.6, p < 0.0001); significant interaction (F_9,467760_= 2.29, p = 0.0144)). This deficit was selective to n - 1 (multiple comparison corrected post-hoc n - 1: t_467760_ = 3.09, p = 0.021). There was no effect on the Moving Average (Figure 6F). Interestingly, complex LMEs revealed a more specific deficit in M2-DMS lesions relative to broad M2 populations; M2-DMS lesions reversed the contribution of a checking HE between press n and press n - 1, such that checking was now detrimental in the use of prior duration information to guide performance (Figure 6G; t_47352_ = 3.10, p = 0.002). However, no other terms differed between Sham and Lesion groups (Table S7).

**Figure 6.**
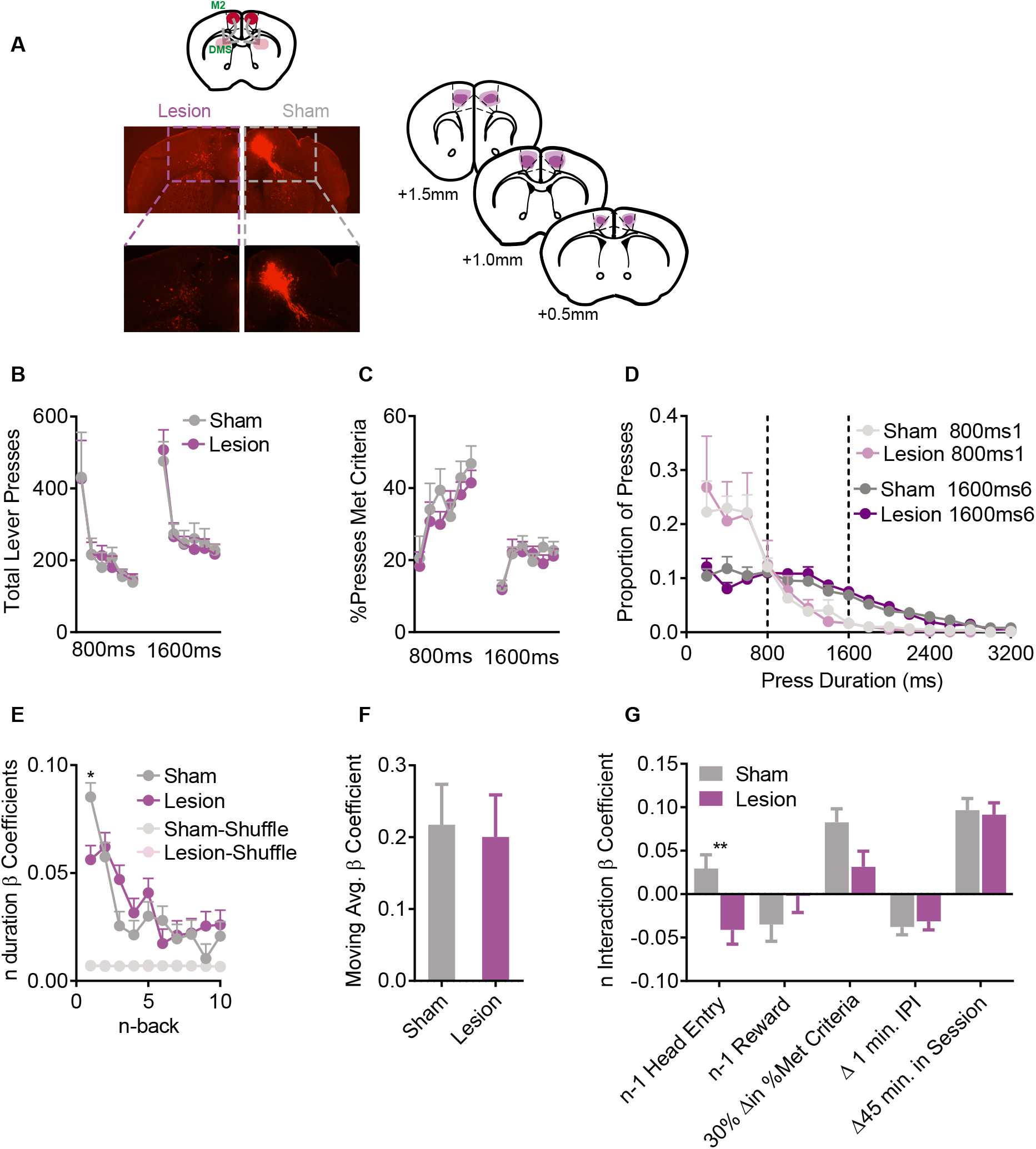
Pretraining M2-DMS lesions impair use of recent experience. (A) (top) Schematic, (bottom) example histology and (right) average/maximal (dark/light shading) spread of projection specific M2-DMS lesion using a Cre-dependent caspase strategy, slice coordinates relative to Bregma. (B) Total Lever Presses across training days. (C) %Presses that met criteria across training days. (D) Histogram of lever press durations on the first and last days of the 800ms and 1600ms criteria (200ms bins). (E) β coefficient from LME models predicting n duration from n-back durations for Actual and order Shuffled data. (F) Moving average β coefficient for Actual and Shuffled data. (G) β coefficients for the interaction terms from the complex LME model. For display purposes we transformed the continuous variables to show relevant changes. B-D are mean+SEM across mice. Shuffled data are the mean+SEM of 1000 order shuffled β coefficients. ** p < 0.01, * p < 0.05. See also Table S7.

The photometry and lesion data together suggest that M2-DMS activity represents and is functionally necessary for recent sequential action (pressing and checking) experience to contribute to the initiation and execution of the current decision. To directly test this hypothesis, we took a behaviorally-dependent, closed-loop optogenetic approach to inhibit M2-DMS neural activity. We used a dual-virus strategy to express an inhibitory opsin (ArchT: n = 5, Figure 7A) that reduced M2-DMS spiking when activated by light (Figure 7B). We targeted inhibition to three different epochs: the initiation of a lever press, during the press itself, and after lever press release. Each manipulation occurred across 6 days of training and only on a subset of lever presses (STAR methods). This allowed us to include additional terms in our LME models to determine if inhibition directly affected press n duration, and/or if inhibition affected the contribution of prior experience (i.e., an interaction between inhibition and n - 1 duration). In addition to this within subject comparison, we also made between subject comparisons to fluorophore control mice (tdTomato: n = 6).

**Figure 7.**
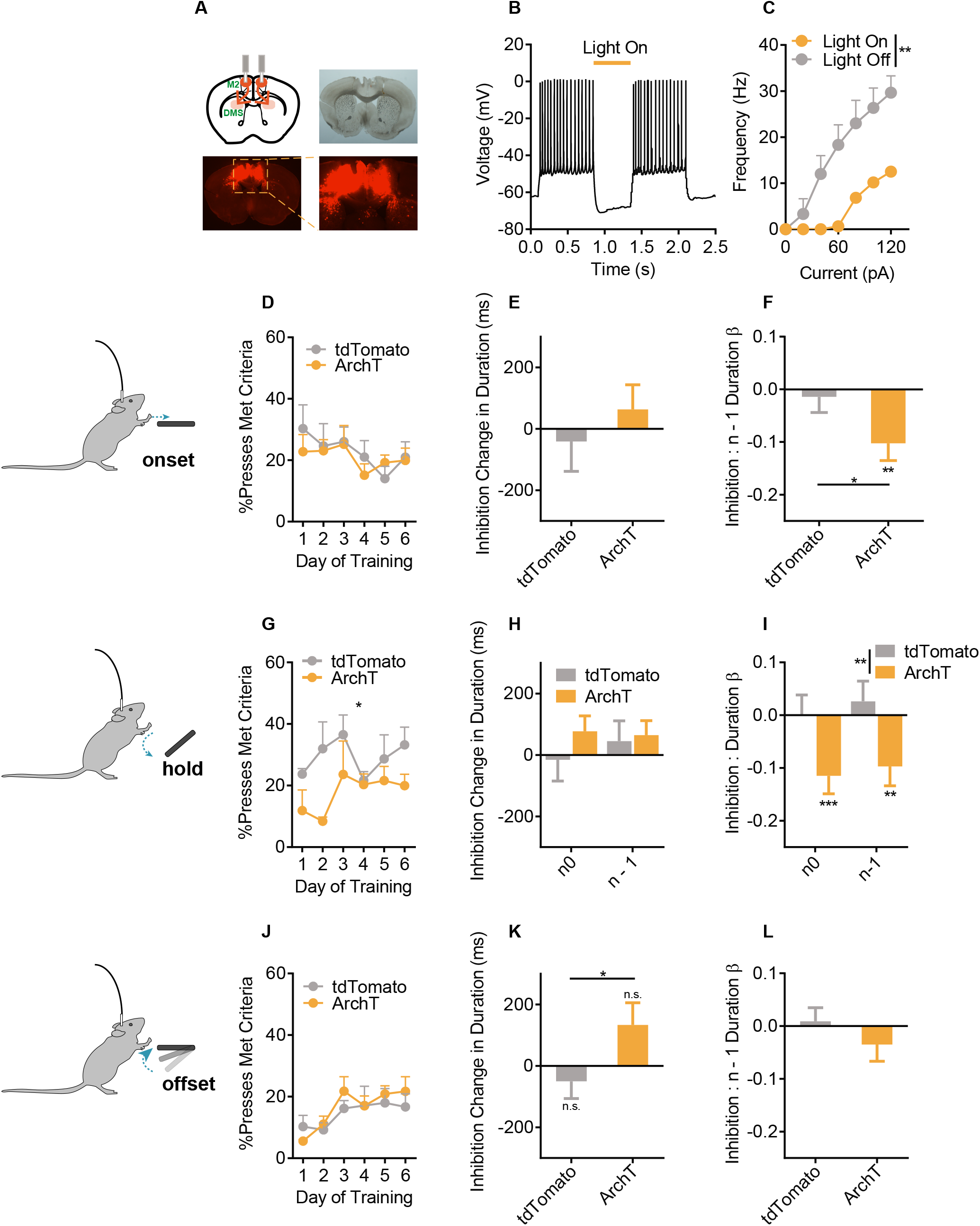
Optogenetic inhibition of M2-DMS projections during execution impairs use of recent experience. (A) (top left) Schematic and example histology of ArchT optogenetic inhibition of M2-DMS projection neurons. (B-C) Slice verification of ArchT-mediated inhibition in M2-DMS projection neurons. ** = 2-way RM ANOVA (Current x Light) interaction: F_6,6_ = 17.0, p = 0.002. (D-F) Pre-onset inhibition. (D) %Presses that met criteria, (E) Main effect of inhibition on duration, and (F) Interaction between inhibition and the contribution of the prior duration. (G-I) As in D-F except for inhibition during the duration of the press. (J-L) As in D-F except for inhibition occurring after press offset. Note that although there is a group difference between ArchT and tdTomato mice in (K), the main effect itself is n.s. in the model for both groups. n0 = Light occurred on press n. n - 1 = Light occurred on press n - 1. tdTomato = tdTomato expressing control mice. ArchT = ArchT expressing mice. In all LME graphs, * without any lines indicate significant terms via F-test on the model (within group comparison), while * with a line indicate significant, between group differences. *** p < 0.001, ** p < 0.01, * p < 0.05. n.s. = Not significant. Data are mean+SEM.

In order to target inhibition prior to press onset, mice were tracked using an overhead camera and light (1s, continuous) was triggered 50% of the time when mice entered a zone centered on the lever. We did not find any effect of pre-onset M2-DMS inhibition on overall performance (Figure 7D), nor any effect on press duration itself (i.e., no main effect of inhibition, Figure 7E). Within the ArchT group, inhibition prior to lever pressing did induce a significant negative interaction with n - 1 duration (F_1,2718_ = 10.6, p = 0.001), and a significant difference with the tdTomato group (Figure 7F; t_6645_ = 2.04, p = 0.042). However, as this inhibition continued for 1s, it may have persisted during lever pressing itself. Indeed, there was no longer an effect of inhibition even within the ArchT group (F_1,534_ = 0.45, p = 0.50) when we limited our analysis to light inhibition that did not spillover into the lever press. Thus, it seems likely that the marginal effect of pre-onset inhibition is due to spillover of inhibition to the lever press itself as opposed to inhibition that occurred prior to press onset.

To directly address this question, we next tied inhibition to lever pressing itself. We inhibited during the full duration of every 7th lever press. Such inhibition did reduce the efficacy of lever pressing (Figure 7G; 2-way RM-ANOVA (Opsin/Fluorophore X Day), main effect only of Opsin group (F_1,9_ = 7.59, p = 0.0223)). Again there was no main effect of inhibition on press duration itself, and this was true both whether inhibition occurred during press n or on press n - 1 (Figure 7H). However, M2-DMS inhibition during press n prevented the use of n - 1 duration information from guiding the current action (Figure 7I). Further, inhibition during press n also prevented the experiential information gained during the execution of press n from informing the next press (F-test within ArchT model, n0: F_1,6110_ = 11.2, p = 0.0008; n - 1: F_1,6110_ = 6.91, p = 0.009. Group comparison between ArchT/tdTomato 2-way ANOVA (Opsin x n-back) main effect only of Opsin group F_1,23602_ = 10.5, p = 0.0012). This suggests that M2-DMS activity during the press itself is not important for controlling the duration of the current lever press *per se*. Instead, this activity contributes to using recent experiential information to execute current actions, and this abrupt disruption impaired task performance. In support of this, when we targeted inhibition to press offset (1s of light after press release), there was neither a direct effect of inhibition on subsequent press durations, nor an interaction with the use of recent experience, nor any effect on performance (Figures 7J-L). There was also no interaction with the moving average term at any inhibition time point. The lack of a direct effect of inhibition on press duration suggests the deficit in use of recent experience was not due to a non-specific motor effect. The lack of any inhibition effect prior to or after execution of the lever press also suggests M2-DMS activity does not represent a form of working memory, but instead supports use of prior experience to inform action execution.

## Discussion

There is a growing concern that neuroscience investigations into decision-making are “missing the forest for the trees”, or *vice versa*. Investigations into the nature of decision-making that isolate specific task-based computations or focus on summary statistics such as accuracy have been indispensable in providing information about both the tree and forest, respectively. However, the present data suggests the need to account for the mesoscopic context experiential information provides in order to link these levels of analysis, akin to understanding the intertwined communication among trees in a forest (Gorzelak et al., 2015). Here, sources of experiential information often treated as task-irrelevant determined whether and to what degree recent experience-information influenced adaptive behavior to support ongoing decision-making. Experiential information influenced the neural correlates of decision-making and determined circuit recruitment and contribution in mice, suggesting such information should not be ignored. By using this approach, we show that M2 and M2-DMS circuits use broad experiential information to instigate exploratory or recent experience-based responding.

Classic temporal difference or reinforcement learning models (Rescorla and Wagner, 1972; Sutton and Barto, 1998) emphasize the role played by responses and outcomes. By using an unconstrained task with a continuous decision variable, we found mice do not drastically shift their strategy solely on their sequence of actions, nor based on whether their action earned a reward. Rather, experiential variables such as checking and the passage of time strongly influenced the behavior and may serve to arbitrate between strategies. Behaviorally the latter could function to bias exploratory responding when the preceding action is more distant in time and hence, when the environment, and/or its neural representation may have changed. On the other hand, the former suggests information seeking itself increases use of experience-based strategies, perhaps as a result of providing definitive feedback. As multiple behavioral controllers can be used to make seemingly similar decisions (Balleine, 2019), experiential information may be used to bias strategy-level recruitment for instance, by adjusting the relative degree of exploration (Figure 3) or the relative similarity between adjacent decisions (Figures 6–7). This bias in recruitment strategy may arise through experiential modulation of associated neural activity (Schreiner and Gremel, 2018), perhaps by setting the “gain” on behavioral strategies (Johnson et al., 2016).

### Integration and implementation of subjective experience in M2 and M2-DMS circuits

We found a robust representation of subjective experience in broader M2 population activity. That lesions both increase the similarity between adjacent presses and reduce the integration of non-action-outcome information (including Checking, IPI, Time, and overall performance) suggests M2 lesions may render mice relatively inflexible and left to rely more on a simple repetition-based strategy. Though it must be noted that the M2 Sham comparison group displays a low n - 1 β coefficient, perhaps inflating the overall effect of M2 Lesions. However, the increased relationship is still present when comparisons are made to another sham group (M2-DMS Shams; t_56426_ = 2.22, p = 0.026), with a similar loss of interactions with other experiential information. Collectively, our results extend previous studies implicating M2 populations in goal-directed or model-based decision-making (Gremel and Costa, 2013) by providing novel insight into precisely *how* this effect is achieved. Namely, by nullifying the contribution of subjective experience in arbitrating between decision-strategies, animals with M2 lesions rely on repetition-based strategies.

While M2 is important for a broader representation of experiential information, in a subset of M2 projections neurons (M2-DMS) we see a more limited contribution of information used to guide ongoing actions. Converging Ca^2+^ activity, lesion, and optogenetic inhibition studies implicate M2-DMS projections specifically in contributing recent action information to ongoing actions. The reduction in the use of recent experience only when optogenetic inhibition occurred *during* the press suggests that M2-DMS activity may serve as an experience-based guide or reference for ongoing actions. This raises the hypothesis that M2-DMS may function as a comparator for template or pattern matching during action performance, analogous to the pattern matching seen in avian vocal learning, and hypothesized to be implemented in premotor regions (Mooney, 2009). In M2-DMS lesioned mice, an intermediate behavior of checking in between lever presses reduced the reliance of the current action on the prior, providing some evidence that M2-DMS function is necessary to link and/or compare recent action experience as has been suggested by work examining sequence learning and initiation (Rothwell et al., 2015). Future studies investigating M2-DMS function at the single neuron level could reveal important insights into precisely how this is instantiated in the brain, and if there is an “embodied engram” of recent actions, or a comparator function in M2-DMS projection neurons.

### Conclusion

Rarely is behavior in the natural world so neatly constrained as in many laboratory tasks; thus it seems likely that animals have adapted to use diverse sources of information to guide their behavior. The brain should therefore be sensitive to this information, yet several recent studies have demonstrated remarkably widespread coding of variables in the brain (Allen et al., 2017; Steinmetz et al., 2019). Perhaps this apparent distributed coding is the consequence of attributing relatively static measurements of behavior and human-derived constructs to large neural populations. That there is a wealth of information available to animals and many neural circuits to support decision-making, raises the hypothesis that specific aspects of experiential information may modulate neural function differentially depending on the circuit and the computation. Indeed, we find that M2 is sensitive to many aspects of this experiential information, but examination of a discrete output population (M2-DMS) showed a more selective representation and functional role. Investigations into circuit, synaptic, and molecular mechanisms controlling how subjective experience modulates decision-making will likely be fruitful, akin to increased understanding of arousal modulation of sensory processing (Shimaoka et al., 2018).

Repetitive decision-making is found across many disease states including substance use disorders and Obsessive Compulsive Disorder. M2’s potential human homologues - the pre-supplementary/supplementary motor areas - are accessible to region-specific treatments such as TMS, which have shown promise in disease treatment (Gomes et al., 2012; Hawken et al., 2016; Mantovani et al., 2013). Here we establish that M2-DMS is involved in implementing repetitive or recent-experience-based decisions. This raises the hypothesis that M2-DMS dysfunction may lead to decisions that are inappropriately or excessively repetitive (Corbit et al., 2019). Incorporating subjective experience into the examination of disease-induced brain function during decision-making may increase the likelihood of obtaining enduring findings relevant to the clinical treatment of disease.

## Acknowledgements

This work was funded by F31AA027439 (D.C.S.), DGE-1650112 (C.C.), F32AA026776 (R.R.), R01AA026077 (C.M.G.), and a Whitehall Foundation Award (C.M.G).

## Author Contributions

D.C.S.: Conceptualization, Formal analysis, Funding acquisition, Investigation, Methodology, Visualization, Writing - original draft, Writing - review and editing. C.C.: Formal analysis and Writing - review and editing. R.R.: Investigation and Writing - review and editing. C.M.G.: Conceptualization, Supervision, Funding acquisition, Project administration, Visualization, Writing - original draft, Writing - review and editing.

## STAR Methods

### Resource Availability

#### Lead Contact

Further information and requests for resources and reagents should be directed to the lead contact, Christina M. Gremel.

#### Materials Availability

This study did not generate any new reagents.

#### Data and Code Availability

- The data reported in this paper will be shared by the lead contact upon request.
- The code used to analyze the data from this study is available at: https://github.com/gremellab/Hold-Down-Behavior-GCAMP-Opto-analysis
- Any additional information required to reanalyze the data reported in this paper is available from the lead contact upon request.

### Experimental Model and Subject Details

Similar numbers of male and female C57BL/6J mice (>7 weeks/50 PND) (The Jackson Laboratory, Bar Harbour, ME) were used for experiments. Exploratory analyses for sex differences in the behavioral cohort revealed no differences, and thus we collapsed across sex. All procedures were conducted during the light period and mice had free access to water throughout the experiment. Mice were housed 2–4 per cage on a 14:10 light:dark cycle. Mice were at least 6 weeks of age prior to surgery. Mice were food restricted to 85-90% of their baseline weight 2 days prior to the start of behavioral procedures, and were fed 1–4 hours after the daily training. All experiments were approved by the University of California San Diego Institutional Animal Care and Use Committee and were carried out in accordance with the National Institutes of Health (NIH) “Principles of Laboratory Care”.

### Methods Details

#### Behavioral Procedures

Mice were trained once per day in operant chambers in sound attenuating boxes (Med-Associates, St Albans, VT) in which they pressed a lever (left or right of the food magazine, counterbalanced for location) for an outcome of regular ‘chow’ pellets (20 mg pellet per reinforcer, Bio-Serv formula F0071). Each training session commenced with an illumination of the house light and lever extension and ended after either 60 reinforcers were earned or 90 minutes had elapsed, with the lever retracting and the house light turning off.

On the first day, mice were trained to approach the food magazine to retrieve the pellet outcome (no lever present) on a random time (RT) schedule, with a reinforcer delivered on average every 120 seconds for a total of 60 minutes. Next, mice were trained on a continuous ratio schedule of reinforcement (CRF) across 3 days, where every lever press was reinforced (no duration requirement), with the total possible number of reinforcers increasing (CRF10, 30, and 60).

Following CRF pretraining, the hold down task was introduced. We instituted a duration requirement on lever pressing: animals had to press and hold down the lever for >800ms in order to earn food reward (delivered immediately after press release). Importantly, there were no cues, no timeout period, nor any discrete trials; the lever was always available to mice, until they completed their session. Mice were trained at the >800ms criterion for 6 days, followed by at least 6 days of training at the >1600ms criterion. During all days, timestamps for lever press onset, lever press offset, the onset and offset of headentry into the food magazine, and the delivery of reward were recorded. From this timing information, we were able to calculate durations of lever presses and headentries. Of note, use of Med Associates introduced a 20 ms limit on our time resolution.

##### Outcome Devaluation

In the behavioral mice (Figure 1 and Figure S1, n = 12 total, n = 7 female and n = 5 male), after 8 days of training at >1600ms, we performed outcome specific satiety. Devaluation procedures occurred across two days. In brief, on the valued day, mice had *ad libitum* access to an outcome previously experienced in the home cage for 1 hour before being placed in the training context for a 5 minute, non-reinforced test session. On the devalued day, mice were given 1 hour of *ad libitum* access to the outcome previously earned by lever press, and then underwent a 5 minute, non-reinforced test session in the training context. One mouse consumed less than .1g of the valued outcome during pretraining exposure and was excluded from all devaluation analyses (giving final n = 11). The order of revaluation day was counterbalanced across mice.

##### Probabilistic Reward

A naive group of mice (n = 15, n = 4 female and n = 11 male) were trained for 6 days on >800 ms, followed by 8 days at > 1600ms criteria, and then switched to probabilistic reward, where only a percentage of presses that exceeded the duration criterion were rewarded on a random ratio schedule. These animals were separated into three different probabilistic reward groups: 25%, 50%, and 75% (n = 5 each group) and trained for a further 3 days under the assigned probabilistic schedule.

#### Surgical Procedures

All viral vectors were obtained from the UNC Viral Vector Core (Chapel Hill, NC) or Addgene (Wateron, MA). Mice were anaesthetized with isoflurane (1–2%) and intracranial injections were performed via Hamilton syringe (Reno, NV) targeted at a relatively posterior portion of M2 (from Bregma: AP +1.0mm, L ±0.5mm and V −1.2mm, −1.4mm from the skull), and/or DMS (from Bregma: AP +1.0mm, L ±1.65mm and V −3.0mm, −3.2mm from the skull). Syringes were left in place for five minutes after each injection to allow for diffusion, and all viruses or drugs were infused at a rate of 100nl/min. Mice were given at least two weeks to allow for recovery and viral expression before the start of experimental procedures (at least four weeks for all M2-DMS manipulations). After behavioral testing was concluded, mice were euthanized and brains were extracted and fixed in 4% paraformaldehyde. Localization and spread of viral expression was assessed in 50-100 μm thick brain slices using fluorescent microscopy (Olympus MVX10, Shinjuku, Japan).

For M2 lesions, n = 12 Lesion mice were bilaterally injected with ibotenic acid (10mg/ml, ThermoFisher), while n = 12 Sham lesion mice were injected with vehicle (saline) at M2 (2 injections of 120nl at V −1.4mm and −1.2mm from the skull in each hemisphere). In order to assess excitotoxic lesion presence and spread, brains were sliced at 50um thick, and incubated with propidium iodide (1:10000 in 1xPBS, Chemodex: P0023) and Isolectin-GS IB_4_ Alexa Fluor 488 Conjugate (20:10000, ThermoFisher: l21411), a marker of microglial cells which are recruited via lesions (Lünemann et al., 2006). Brain slices were incubated for 1hr, followed by 3 x 10min washes. 4 Sham mice were excluded due to technical difficulties during training, and 2 Lesion mice were excluded due to histology, giving final n’s of 10 Lesion (n = 6 female, n = 4 male) and 8 Sham (n = 4 female, n = 4 male) mice.

For M2 GCaMP experiments, n = 8 mice (n = 4 female, n = 4 male) were injected (2 injections of 200nl at V −1.4mm and 1.2mm from the skull) with rAAVDJ/PAAV-CaMKIIa-GCaMP6s to express GCaMP6s under control of the Ca^2+^ calmodulin dependent protein kinase IIα (CamKIIα) promoter and implanted with an optical fiber unilaterally in M2.

For M2-DMS GCaMP experiments, n = 8 mice were unilaterally injected with a viral vector expressing Cre recombinase (AAV5/Ef1a-Cre-WPRE) in DMS (2 injection depths: V −3.0mm and −2.8mm from the skull, 250nl each), and were injected with a viral vector expressing a Cre-dependent GCaMP6s (pAAV.CAG.FLEX.GCaMP6s.WPRE.SV40 (Addgene: 100842); 2 injection depths: V: −1.4mm, and −1.2mm from the skull, 200nl each) followed by fiber implantation in ipsilateral M2. One mouse was excluded due to histology (n = 4 female, n = 3 male).

For M2-DMS lesion, n = 8 Lesion (n = 4 female, n = 4 male) and n = 8 Sham (n = 4 female, n = 4 male) mice were bilaterally injected with 200nl of a viral vector expressing CamKIIα-Cre in DMS (rAAV5/CamKII-GFP-Cre; 2 injection depths: V: −3.1mm and −2.9mm from skull, 200nl each). Lesion and Sham mice were also injected with a viral vector expressing Cre-dependent tdTomato in M2 (rAAV5/Flex-tdTomato; 100nl at V −1.4mm from the skull). Lesion mice additionally received a viral vector expressing a Cre-dependent caspase virus in M2 to induce apoptosis of M2-DMS projections (rAAV5/AAV-Flex-taCasP3-TEVP; 2 injection depths: V −1.4mm and −1.2mm from the skull, 200nl each).

For M2-DMS optogenetic inhibition experiments, n = 8 Archaerhodopsin (ArchT) and n = 8 tdTomato mice were bilaterally injected with a viral vector expressing CamKIIα-Cre in DMS (rAAV5/CamKII-GFP-Cre; 2 injection depths: V −3.1mm and −2.9mm from the skull, 200nl each). Following exclusion for viral expression or low levels of behavior, there were n = 5 ArchT mice (n = 3 male, n = 2 female), and n = 6 tdTomato control mice (n = 3 male, n = 3 female). Due to the proximity of bilateral M2 at this posterior portion (~1.0mm) for ferrule implantation, we injected virus and implanted fibers at a 10° angle, and adjusted the M2 coordinates accordingly. Experimental ArchT mice received a viral vector expressing a Cre-dependent inhibitory opsin (rAAV5/Flex-ArchT-tdTomato), while fluorophore control mice received a viral vector expressing Cre-dependent fluorophore only (rAAV5/Flex-tdTomato), in both cases receiving the same injection volume (300nl at V −1.42mm from the skull), with bilateral M2 fibers implanted at V −1.37mm from the skull.

#### Fiber Photometry

Animals underwent pre-training as described above, but received one additional day of CRF training during which animals were first hooked up to 400 um optical fiber tethers with ferrule to ferrule connectivity. A 470nm LED (Thorlabs, Newton, NJ) was used for excitation of GCaMP6s (< 70 μW/mm2), and fluorescence emissions were collected with a bifurcated fiber (Thorlabs, Newton, NJ) which allowed for simultaneous, independent recordings of two mice. We imaged the dual fiber core using a 4x objective (Olympus, Shinjuku, Japan) focused onto a CMOS camera (FLIR Systems, Wilsonville, OR). Regions of interest (i.e., the fiber cores) were selected using Bonsai software (Lopes et al., 2015) to acquire fluorescence intensity signals (at a rate of 20Hz). Bonsai software simultaneously collected analog behavioral data and timestamps for lever presses, head entries, and reinforcer delivery sent via TTL Med-PC pulses using microprocessors (Arduino Duo, from Arduino, Sumerville, MA) with custom code. Photometry and behavioral data were imported into Matlab (Mathworks Inc., Natick, MA) for analysis using custom scripts. To account for decay across the session (photobleaching), we fit the fluorescence intensity signal to a double exponential. To check for bad coupling of the fiber to the ferrule, or low expression each session we calculated the 97.5 percentile of dF/F and ensured that there was at least a 1% change, sessions failing to meet this criterion were excluded from analyses (Markowitz et al., 2018), and also excluded sessions with visual anomalies in the session long traces (e.g., a sudden, sustained decrease in activity partway through the session that could indicate fiber decoupling). We used the mean and standard deviation during a baseline period −15s to −5s prior to lever pressing to z-score press-related activity (i.e., from −5s prior to onset up to 5s post offset). To compare Met and Fail lever presses, we performed running permutation tests, requiring that at least 4 adjacent samples were significantly different from one another to control for fluctuations in the data (functions implemented in Matlab from Jean-Richard-dit-Bressel et al., 2020). We smoothed Ca^2+^ activity data using a 10 sample (or 5 sample for interpolated activity) long Gaussian filter for display purposes only.

#### Optogenetic Inhibition

For bilateral light delivery, Arduino Duos with custom code were used to receive TTL pulses from Med-PC operant chambers and trigger onset of 2 LEDs (595nm, Thorlabs) coupled to 200um sheathed fiber optic cable with ferrule to ferrule connectivity (>= 1mW at ferrule tip). We used 595nm light as this has been shown to optimally activate ArchT while avoiding non-specific effects (Setsuie et al., 2020). We used several different protocols to target the closed-loop inhibition to different task epochs. Inhibition during the duration of the lever press occurred across the 6 >800ms training days, with light delivery (continuous, not pulsed) tied to the lever pressing itself. As we observed a decaying relationship between n-back press durations and n - 0 press duration, every 7th lever press triggered light delivery, which persisted for the duration of the lever press (with a time resolution of 20ms for light offset). During days 1-6 of the >1600ms training, we instead tied light delivery to press *offset*, again, on every 7th lever press. Thus, after every 7th lever press, mice were given 1s of light. Finally, after undergoing 4 days of baseline training without any light inhibition (though while still being hooked up to fibers), we shifted to inhibiting *prior* to press onset for 6 days. In order to achieve this, animals were recorded with an overhead camera (1080p wide angle webcam, Logitech) and tracked in real time using Bonsai software. We individually defined regions of interest centered on the lever (approximately twice the width and length of the lever itself). 50% of entrances into this region generated a TTL pulse to turn on the LEDs, which remained on for 1s.

#### ArchT Slice Validation

Coronal slices (250 μm thick) containing M2 were prepared using a Pelco easiSlicer (Ted Pella Inc, Redding, CA). Mice were anesthetized by inhalation of isoflurane and brains were rapidly removed and placed in 4°C oxygenated ACSF containing the following (in mM): 210 sucrose, 26.2 NaHCO_3_, 1 NaH_2_PO_4_, 2.5 KCl, 11 dextrose, bubbled with 95% O_2_/5% CO_2_. Slices were transferred to an ACSF solution for incubation containing the following (in mM): 120 NaCl, 25 NaHCO_3_, 1.23 NaH_2_PO_4_, 3.3 KCl, 2.4 MgCl_2_, 1.8 CaCl_2_, 10 dextrose. Slices were continuously bubbled with 95% O_2_/5% CO_2_ at pH 7.4, 32°C and were maintained in this solution for at least 60 min prior to recording.

Whole-cell current clamp recordings were made in pyramidal cells of M2. Pyramidal cells that expressed ArchT were identified by the fluorescent tdTomato label using an Olympus BX51WI microscope mounted on a vibration isolation table and a high-power LED (LED4D067, Thorlabs). Recordings were made in ACSF containing (in mM): 120 NaCl, 25 NaHCO_3_, 1.23 NaH_2_PO_4_, 3.3 KCl, 0.9 MgCl_2_, 2.0 CaCl_2_, and 10 dextrose, bubbled with 95% O_2_/5% CO_2_. ACSF was continuously perfused at a rate of 2.0 mL/min and maintained at a temperature of 32°C. Picrotoxin (50 μM) was included in the recording ACSF to block GABAA receptor-mediated synaptic currents. Recording electrodes (thin-wall glass, WPI Instruments) were made using a PC-10 puller (Narishige International, Amityville, NY) to yield resistances between 3–6 MΩ. Electrodes were filled with (in mM): 135 KMeSO_4_, 12 NaCl, 0.5 EGTA, 10 HEPES, 2 Mg-ATP, 0.3 Tris-GTP, 260–270 mOsm (pH 7.3). Access resistance was monitored throughout the experiments. Cells in which access resistance varied more than 20% were not included in the analysis.

Recordings were made using a MultiClamp 700B amplifier (Molecular Devices, Union City, CA), filtered at 2 kHz, digitized at 10 kHz with Instrutech ITC-18 (HEKA Instruments, Bellmore, NY), and displayed and saved using AxographX (Axograph, Sydney, Australia). A series of fixed current injections (20 pA increments from 0 to 120 pA) were used to elicit action potential firing and the number of spikes were counted at each current step. For verification of ArchT function, ArchT was optically stimulated using 590nm light, delivered via field illumination using a high-power LED (LED4D067, Thorlabs). Optical stimulation was done under constant illumination for 1s during current injections.

#### Data Analysis

##### Linear Mixed Effects Models of Behavior

We built simple Linear Mixed Effects (LME) models to model the relationship between the duration of lever press n and n-back (n - 1 through n - 10) lever press durations. We included random intercept terms for mouse and day of training to account for the repeated structure of our data. To determine how far back a significant relationship existed between press n and any particular n-back press, we shuffled the order of a particular n-back (e.g., only n - 3) 1000 times and compared the shuffled distribution of beta coefficients to the actual value via permutation test. Of note, we are shuffling here within individual mouse/sessions, thus preserving the overall statistics of the data, and shuffling only the order in which a specific type of event occurred.

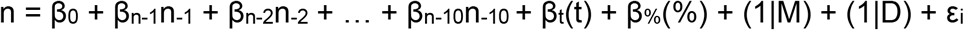

Where n is the current lever press duration, n - 1 through n - 10 are the previous 1 through 10 lever press durations and β_x_ is the linear regression coefficient for term x (β_0_ being the intercept term). We also included covariates of time in session (t) and the percentage of presses that met criteria (%). We included random intercept terms for both mouse (M) and day (D).

In order to determine which other experiential variables affect lever press n duration, we also built more complex LME models that included additional variables. To select variables for this model, we created a “full” model that included n-back durations up to n - 6 (as that is as far back as we see a consistent difference from shuffled data in the simple models), and then main effects of other variables and their interactions with n-back durations, also up to n - 6 (e.g., a binary for if mice made a checking headentry after the previous lever press). We individually removed terms from this full model, and compared Bayesian Information Criterion (BIC) to assess if adding a term improved the model. If any term did not improve the model, we removed it, and also removed any further n-back examples of it. However, we kept main effect terms in the model if the interactions were significant, and kept all the same interaction terms for n - 1 and the moving average term to be able to directly compare how various events might differentially affect the contribution of press n - 1 versus the moving average. To ensure that terms in this reduced model did not improve the model due to overall correlations across days or mice, we also compared beta coefficients from the actual data to those obtained from 1000 order shuffled datasets, where we individually permuted a given term within individual mouse/sessions. This analysis conducted on our “reduced” model agreed with the BIC selection for terms that improved the model. We were ultimately left with the model in Table S3 (see also Table S2 for a description of the terms), signified by the equation below.

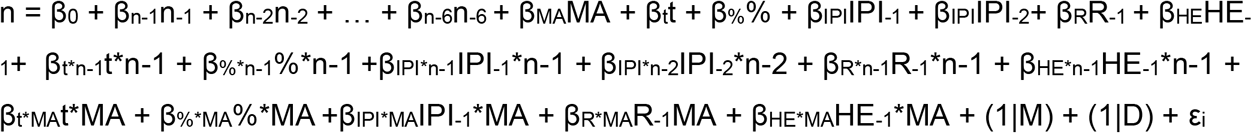

Where β_x_ represents the linear regression coefficient for a given term. This model has the same terms as the simple model, though only back to n - 6 durations, as that is as far back as there is a reliable difference to shuffled data. In addition, there is the MA term which is a moving average from presses n - 7 through n - 60 (length selected via BIC using different window lengths). Additionally, we have main effects of time in session (t, in ms), the percentage of presses that met criteria (%), inter-press interval (IPI in ms, for both time in between press n and press n - 1 (IPI_−1_), and between press n and press n - 2 (IPI_−2_)), outcome of press n - 1 (R_−1_: binary where 0 is no reward and 1 is reward), and headentry between press n - 1 and press n (HE_−1_: binary where 0 is no headentry and 1 is headentry). Again, we have random intercept terms for mouse (M) and day of training (D). We also included interaction terms between the n - 1 duration term and: t, %, IPI, R_n-1_, and HE_n-1_. These interaction terms are specified with the general format of β_x*n-1_x*n-1 where x represents an individual interaction term (e.g., for time in session t interacting with n - 1 duration: β_t*n-1_t*n-1).These same interaction terms were included with the moving average term (MA, of the general format β_x*MA_x*MA) in order to see if very recent experience (n - 1) and long-term experience (MA) were differentially influenced by variables such as time. Interestingly, when examining further n-back interactions, only the interaction between IPI_n-2_ and n - 2 duration survived the BIC selection process, indicating that individual further n-backs were less open to modification by these variables.

In the probabilistic reward experiment, we added a trinary term for if a lever press was unsuccessful (0), successful and rewarded (1), or successful and unrewarded (2), and included interactions between this term and n - 1 as well as the MA. Additionally, we ran all three probability groups together in the model and included indicator variables for which group (25%, 50%, or 75% reward) a mouse belonged to. This allowed us to include a 3-way interaction to determine if the groups differed in how this trinary outcome term interacted with prior press durations (e.g., does the probability of reward influence the presence/magnitude of win-stay behavior?). For the optogenetic inhibition LME models, we included a binary term indicating if a lever press received light delivery (before, during, or after for the three different manipulations) as both a main effect and as an interaction with n - 1 duration and the MA to determine if light reduced the relationship between press n and press n - 1/the MA.

##### Ca^2+^ Activity Linear Mixed Effect Models

For the M2 and M2-DMS Ca^2+^ activity recordings, we built LME models that sought to predict Ca^2+^ activity given behavior. For this, we used data only from the 1600ms training days. First, we built simple LME models that included only current (n) and prior (n-back, up to n - 6) durations to predict activity (calculated as area under the curve) at three different time points: −1s to 0s before press onset, during the lever press itself, and 0s to +1s after press release. For activity during the lever press itself, we used modified Akima interpolation, implemented using Matlab’s interp1 function to get presses of different durations on the same relative scale, and we excluded any lever presses with fewer than 2 samples which would preclude interpolation. We also included terms for prior activity (up to n - 6) to control for autocorrelation in the Ca^2+^ activity signal. We again compared beta coefficients from the actual data to 1000 order shuffled datasets for these simple models.

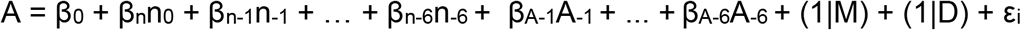

Where A is current Ca^2+^ activity and β_x_ is the regression coefficient for term x. Of note, these models included n duration (n_0_) as a predictor (whereas this was what we predicted in the pure behavioral models). We predicted A given both current (n_0_) and prior (n - 1, up to n - 6) press durations, included prior Ca^2+^ activity (A - 1, up to A - 6) as a covariate, and included random intercepts of mouse (M) and training day (D).

Additionally, we built more complex LME models to predict Ca^2+^ activity data. For these, we used the complex behavioral model above for the predictors, as we were interested in seeing if these variables - which we know influence the behavior - are also represented in Ca^2+^ activity, and also still included prior Ca^2+^ activity data to control for autocorrelation in the Ca^2+^ data. This took the form of the following equation, using the same variables as the preceding equations.

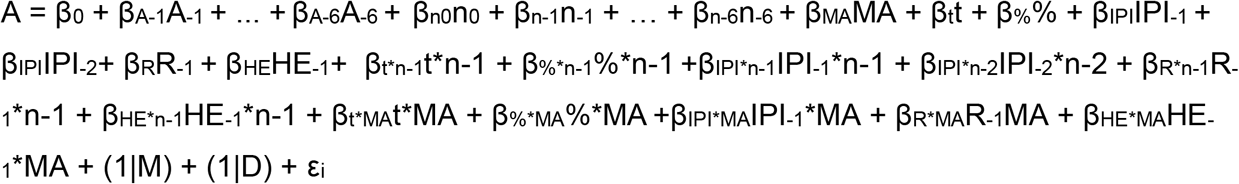

When trying to predict activity after lever press offset, we also included a binary term for outcome on lever press n (R_0_ i.e., the lever press that was just completed with 0 being no reward and 1 being reward). We did not include this term at the other time points since it would introduce a “post diction” confound (i.e., including a term for the outcome of a press before the press even occurred at onset). For the same reason, we did not include interactions with the n_0_ variable.

### Quantification and Statistical Analysis

All analyses were two-tailed with α = 0.05 as a threshold for significance. For analyzing coarse behavioral measurements (e.g., Total Lever Presses) one-way or two-way RM ANOVAs were used, with Greenhouse-Geisser correction for one-way ANOVA and Bonferroni corrections for post-hoc multiple comparisons unless otherwise noted. We used the RMcorr package (Bakdash and Marusich, 2017) implemented in R (R core team, 2014) to calculate a repeated measures correlation between individual model fit and mouse performance to account for the repeated nature of this data (sampling individual mice across days). We used Matlab’s cusum function to get the upper cumulative sum in Figures 1I-J, using 2SD as the criterion. In our simple LME models, we used permutation tests comparing actual β coefficient values to a distribution of 1000 order shuffled versions of the same variable, and thus the resolution of our permutation p-values was p < 0.001. We excluded presses over 10s in duration from all modeling datasets. For event-aligned Ca^2+^ activity comparing Met vs. Fail lever presses, we used permutation tests that required either 4 (for onset and offset-aligned activity) or 3 (for interpolated activity during the press) consecutive samples passed the threshold for significance. To assess the relationship between Ca^2+^ activity and various aspects of behavior in our complex LME models, we performed F-tests on the individual parameters. For group comparisons (e.g., Sham vs. Lesion) of LME model coefficients, we used t-tests with Benjamini-Hochberg false discovery rate correction (Q = 5%) on all of the model terms shown in Tables S4 and S7. Behavioral data was analyzed using Excel (Microsoft), Matlab (Mathworks), R (R core team, 2014), and Prism (Graphpad).

## References

Adams, C.D., and Dickinson, A. (1981). Instrumental responding following reinforcer devaluation. Q. J. Exp. Psychol. Sect. B 33, 109–121.

Allain, F., Minogianis, E.-A., Roberts, D.C.S., and Samaha, A.-N. (2015). How fast and how often: The pharmacokinetics of drug use are decisive in addiction. Neurosci. Biobehav. Rev. 56, 166–179.

Allen, W.E., Kauvar, I.V., Chen, M.Z., Richman, E.B., Yang, S.J., Chan, K., Gradinaru, V., Deverman, B.E., Luo, L., and Deisseroth, K. (2017). Global Representations of Goal-Directed Behavior in Distinct Cell Types of Mouse Neocortex. Neuron 94, 891–907.e6.

Ariely, D., and Zakay, D. (2001). A timely account of the role of duration in decision making. Acta Psychol. 108, 187–207.

Bakdash, J.Z., and Marusich, L.R. (2017). Repeated Measures Correlation. Front. Psychol. 8.

Balleine, B.W. (2019). The Meaning of Behavior: Discriminating Reflex and Volition in the Brain. Neuron 104, 47–62.

Balleine, B.W., and Dickinson, A. (1998). Goal-directed instrumental action: contingency and incentive learning and their cortical substrates. Neuropharmacology 37, 407–419.

Berridge, K.C., Zhang, J., and Aldridge, J.W. (2008). Computing motivation: Incentive salience boosts of drug or appetite states. Behav. Brain Sci. 31, 440–441.

Bouton, M.E., and Balleine, B.W. (2019). Prediction and control of operant behavior: What you see is not all there is. Behav. Anal. Res. Pract. 19, 202–212.

Busse, L., Ayaz, A., Dhruv, N.T., Katzner, S., Saleem, A.B., Schölvinck, M.L., Zaharia, A.D., and Carandini, M. (2011). The Detection of Visual Contrast in the Behaving Mouse. J. Neurosci. 31, 11351–11361.

Corbit, V.L., Manning, E.E., Gittis, A.H., and Ahmari, S.E. (2019). Strengthened inputs from secondary motor cortex to striatum in a mouse model of compulsive behavior. J. Neurosci. 39, 2965–2975.

Costa, R.M. (2011). A selectionist account of de novo action learning. Curr. Opin. Neurobiol. 21, 579–586.

Cristol, D.A., and Switzer, P.V. (1999). Avian prey-dropping behavior. II. American crows and walnuts. Behav. Ecol. 10, 220–226.

Delevich, K., Okada, N.J., Rahane, A., Zhang, Z., Hall, C.D., and Wilbrecht, L. (2020). Sex and Pubertal Status Influence Dendritic Spine Density on Frontal Corticostriatal Projection Neurons in Mice. Cereb. Cortex 30, 3543–3557.

Dhawale, A.K., Miyamoto, Y.R., Smith, M.A., and Ölveczky, B.P. (2019). Adaptive Regulation of Motor Variability. Curr. Biol. 29, 3551-3562.e7.

Dickinson, A. (1985). Actions and Habits: The Development of Behavioural Autonomy. Philos. Trans. R. Soc. B Biol. Sci. 308, 67–78.

Ebbesen, C.L., Insanally, M.N., Kopec, C.D., Murakami, M., Saiki, A., and Erlich, J.C. (2018). More than Just a “Motor”: Recent Surprises from the Frontal Cortex. J. Neurosci. 38, 9402–9413.

Erlich, J.C., Bialek, M., and Brody, C.D. (2011). A Cortical Substrate for Memory-Guided Orienting in the Rat. Neuron 72, 330–343.

Fan, D., Rossi, M.A., and Yin, H.H. (2012). Mechanisms of Action Selection and Timing in Substantia Nigra Neurons. J. Neurosci. 32, 5534–5548.

Gibbon, J., Church, R.M., and Meck, W.H. (1984). Scalar Timing in Memory. Ann. N. Y. Acad. Sci. 423, 52–77.

Gomes, P.V.O., Brasil-Neto, J.P., Allam, N., and Rodrigues de Souza, E. (2012). A Randomized, Double-Blind Trial of Repetitive Transcranial Magnetic Stimulation in Obsessive-Compulsive Disorder With Three-Month Follow-Up. J. Neuropsychiatry Clin. Neurosci. 24, 437–443.

Gomez-Marin, A., Paton, J.J., Kampff, A.R., Costa, R.M., and Mainen, Z.F. (2014). Big behavioral data: psychology, ethology and the foundations of neuroscience. Nat. Neurosci. 17, 1455–1462.

Gorzelak, M.A., Asay, A.K., Pickles, B.J., and Simard, S.W. (2015). Inter-plant communication through mycorrhizal networks mediates complex adaptive behaviour in plant communities. AoB Plants. 7.

Gremel, C.M., and Costa, R.M. (2013). Premotor cortex is critical for goal-directed actions. Front. Comput. Neurosci. 7.

Hawken, E.R., Dilkov, D., Kaludiev, E., Simek, S., Zhang, F., and Milev, R. (2016). Transcranial Magnetic Stimulation of the Supplementary Motor Area in the Treatment of Obsessive-Compulsive Disorder: A Multi-Site Study. Int. J. Mol. Sci. 17.

Hintiryan, H., Foster, N.N., Bowman, I., Bay, M., Song, M.Y., Gou, L., Yamashita, S., Bienkowski, M.S., Zingg, B., Zhu, M., et al. (2016). The mouse cortico-striatal projectome. Nat. Neurosci. 19, 1100–1114.

Iigaya, K., Fonseca, M.S., Murakami, M., Mainen, Z.F., and Dayan, P. (2018). An effect of serotonergic stimulation on learning rates for rewards apparent after long intertrial intervals. Nat. Commun. 9, 1–10.

Jean-Richard-dit-Bressel, P., Clifford, C.W.G., and McNally, G.P. (2020). Analyzing Event-Related Transients: Confidence Intervals, Permutation Tests, and Consecutive Thresholds. Front. Mol. Neurosci. 13.

Johnson, C.M., Peckler, H., Tai, L.-H., and Wilbrecht, L. (2016). Rule learning enhances structural plasticity of long-range axons in frontal cortex. Nat. Commun. 7, 10785.

Juavinett, A.L., Erlich, J.C., and Churchland, A.K. (2018). Decision-making behaviors: weighing ethology, complexity, and sensorimotor compatibility. Curr. Opin. Neurobiol. 49, 42–50.

Klaus, A., da Silva, J.A., and Costa, R.M. (2019). What, If, and When to Move: Basal Ganglia Circuits and Self-Paced Action Initiation. Annu. Rev. Neurosci. 42, 459-483.

Krakauer, J.W., Ghazanfar, A.A., Gomez-Marin, A., MacIver, M.A., and Poeppel, D. (2017). Neuroscience Needs Behavior: Correcting a Reductionist Bias. Neuron 93, 480–490.

Lak, A., Hueske, E., Hirokawa, J., Masset, P., Ott, T., Urai, A.E., Donner, T.H., Carandini, M., Tonegawa, S., Uchida, N., et al. (2020). Reinforcement biases subsequent perceptual decisions when confidence is low, a widespread behavioral phenomenon. ELife 9, e49834.

Lopes, G., Bonacchi, N., Frazão, J., Neto, J.P., Atallah, B.V., Soares, S., Moreira, L., Matias, S., Itskov, P.M., Correia, P.A., et al. (2015). Bonsai: an event-based framework for processing and controlling data streams. Front. Neuroinformatics 9.

Lünemann, A., Ullrich, O., Diestel, A., Jöns, T., Ninnemann, O., Kovac, A., Pohl, E.E., Hass, R., Nitsch, R., and Hendrix, S. (2006). Macrophage/microglia activation factor expression is restricted to lesion-associated microglial cells after brain trauma. Glia 53, 412–419.

Mantovani, A., Rossi, S., Bassi, B.D., Simpson, H.B., Fallon, B.A., and Lisanby, S.H. (2013). Modulation of motor cortex excitability in obsessive-compulsive disorder: An exploratory study on the relations of neurophysiology measures with clinical outcome. Psychiatry Res. 210, 1026–1032.

Markowitz, J.E., Gillis, W.F., Beron, C.C., Neufeld, S.Q., Robertson, K., Bhagat, N.D., Peterson, R.E., Peterson, E., Hyun, M., Linderman, S.W., et al. (2018). The Striatum Organizes 3D Behavior via Moment-to-Moment Action Selection. Cell 174, 44-58.e17.

McDougle, S.D., Butcher, P.A., Parvin, D.E., Mushtaq, F., Niv, Y., Ivry, R.B., and Taylor, J.A. (2019). Neural Signatures of Prediction Errors in a Decision-Making Task Are Modulated by Action Execution Failures. Curr. Biol. 29, 1606-1613.e5.

Mooney, R. (2009). Neural mechanisms for learned birdsong. Learn. Mem. 16, 655–669.

Murakami, M., Vicente, M.I., Costa, G.M., and Mainen, Z.F. (2014). Neural antecedents of self-initiated actions in secondary motor cortex. Nat. Neurosci. 17, 1574–1582.

Murakami, M., Shteingart, H., Loewenstein, Y., and Mainen, Z.F. (2017). Distinct Sources of Deterministic and Stochastic Components of Action Timing Decisions in Rodent Frontal Cortex. Neuron 94, 908-919.e7.

Pinto, L., Rajan, K., DePasquale, B., Thiberge, S.Y., Tank, D.W., and Brody, C.D. (2019). Task-Dependent Changes in the Large-Scale Dynamics and Necessity of Cortical Regions. Neuron. 104, 810-824.e9.

Pisupati, S., Chartarifsky-Lynn, L., Khanal, A., and Churchland, A.K. (2021). Lapses in perceptual decisions reflect exploration. ELife 10, e55490.

Platt, J.R., Kuch, D.O., and Bitgood, S.C. (1973). Rats’ lever-press durations as psychophysical judgments of time. J. Exp. Anal. Behav. 19, 239–250.

R Core Team (2014). R: A language and environment for statistical computing. R Foundation for Statistical Computing, Vienna, Austria. Available at: http://www.R-project.org/

Remedios, R., Kennedy, A., Zelikowsky, M., Grewe, B.F., Schnitzer, M.J., and Anderson, D.J. (2017). Social behaviour shapes hypothalamic neural ensemble representations of conspecific sex. Nature 550, 388–392.

R.A. Rescorla, A.R. Wagner. (1972). A theory of Pavlovian conditioning: Variations in the effectiveness of reinforcement and nonreinforcement. Classical Conditioning II: Current Research and Theory, 2 (1972), pg 64-99.

Rothwell, P.E., Hayton, S.J., Sun, G.L., Fuccillo, M.V., Lim, B.K., and Malenka, R.C. (2015). Input- and Output-Specific Regulation of Serial Order Performance by Corticostriatal Circuits. Neuron 88, 345–356.

Roy, N.A., Bak, J.H., Akrami, A., Brody, C.D., and Pillow, J.W. (2021). Extracting the dynamics of behavior in sensory decision-making experiments. Neuron 109, 597-610.e6.

Schreiner, D.C., and Gremel, C.M. (2018). Orbital Frontal Cortex Projections to Secondary Motor Cortex Mediate Exploitation of Learned Rules. Sci. Rep. 8.

Schreiner, D.C., Yalcinbas, E.A., and Gremel, C.M. (2021). A push for examining subjective experience in value-based decision-making. Curr. Opin. Behav. Sci. 41, 45–49.

Setsuie, R., Tamura, K., Miyamoto, K., Watanabe, T., Takeda, M., and Miyashita, Y. (2020). Off-Peak 594-nm Light Surpasses On-Peak 532-nm Light in Silencing Distant ArchT-Expressing Neurons In Vivo. iScience 23, 101276.

Shimaoka, D., Harris, K.D., and Carandini, M. (2018). Effects of Arousal on Mouse Sensory Cortex Depend on Modality. Cell Rep. 22, 3160–3167.

Siniscalchi, M.J., Phoumthipphavong, V., Ali, F., Lozano, M., and Kwan, A.C. (2016). Fast and slow transitions in frontal ensemble activity during flexible sensorimotor behavior. Nat. Neurosci. 19, 1234–1242.

Skinner, B.F. (1938). The behavior of organisms: an experimental analysis (Oxford, England: Appleton-Century).

Steinmetz, N.A., Zatka-Haas, P., Carandini, M., and Harris, K.D. (2019). Distributed coding of choice, action and engagement across the mouse brain. Nature 576, 266-273.

R.S. Sutton, A.G. Barto. (1998). Reinforcement Learning: An Introduction. MIT Press, Cambridge, MA (1998).

Tervo, D.G.R., Proskurin, M., Manakov, M., Kabra, M., Vollmer, A., Branson, K., and Karpova, A.Y. (2014). Behavioral Variability through Stochastic Choice and Its Gating by Anterior Cingulate Cortex. Cell 159, 21–32.

Yin, H.H., and Yin, H.H. (2009). The role of the murine motor cortex in action duration and order. Front. Integr. Neurosci. 3, 23.

Yoo, S.B.M., Hayden, B.Y., and Pearson, J.M. (2021). Continuous decisions. Philos. Trans. R. Soc. B Biol. Sci. 376, 20190664.

