## Supplemental Figures S1-2 and Tables S1-7 for "Mice are not automatons; subjective experience in premotor circuits guides behavior"

### Figure S1

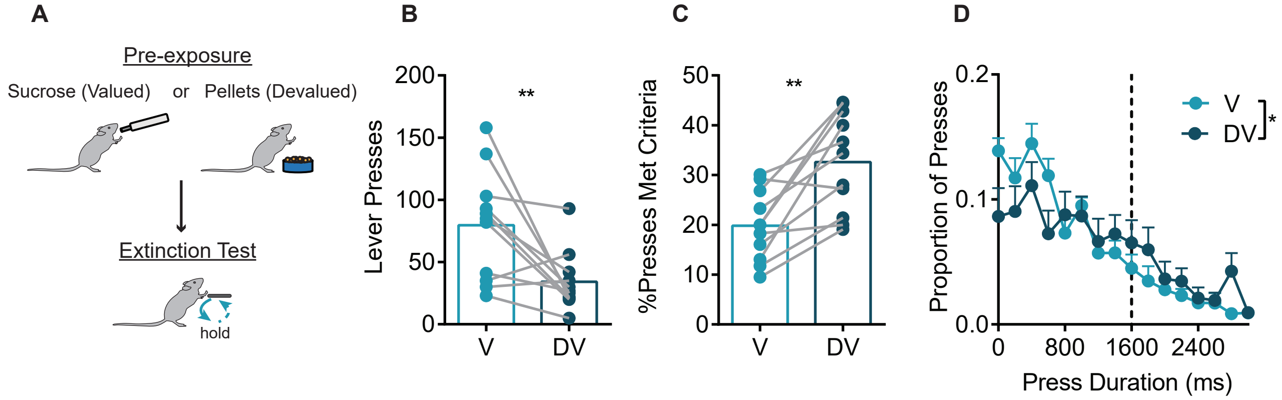


**Figure S1**. *Outcome value has differential control over action selection and execution.* Related to Figure 1. (A) Schematic of the outcome devaluation procedure. In a within subjects design, mice received 1 hr of pre-exposure to either sucrose (Valued, V) or pellets (Devalued, DV), followed by a 5 minute extinction test across 2 days (order counterbalanced). (B) Total Lever Presses on V and DV days. Paired t-test, $t_{10} = 3.09$, $p = 0.012$. (C) %Presses that met criteria on V and DV days. Paired t-test, $t_{10} = 4.55$, $p = 0.0011$. (D) Histogram of press durations on V and DV days (200ms bins). 2-way RM ANOVA, main effect of Duration Bin, $F_{15,150} = 12.1$, $p < 0.0001$ and an interaction (Duration Bin x V/DV) $F_{15,150} = 2.19$, $p = 0.009$. Data in (D) are mean+SEM across mice, bars in (B-C) are mean. ** $p < 0.01$, * $p < 0.05$.

### Figure S2

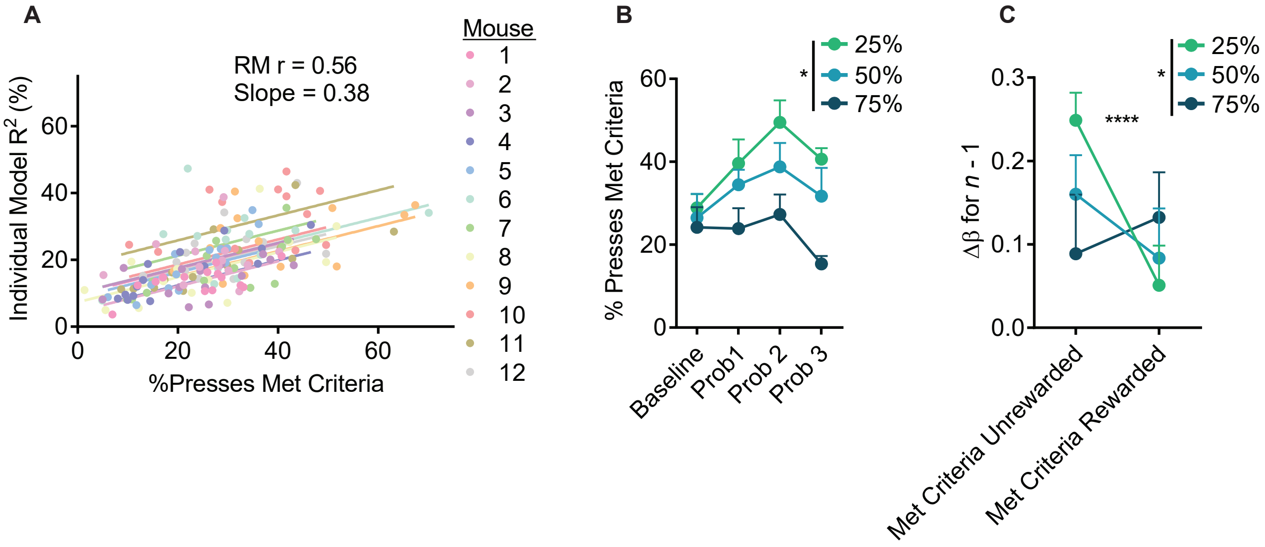


**Figure S2**. *Probabilistic reward induces win-stay behavior and use of experience correlates with task performance.* Related to Figure 2. (A) Repeated measures correlation between task performance (%Presses Met Criteria) and model fit ($R^2$) when building LMEs using individual mouse/session data. Intercept was allowed to vary across mice while using a common slope. (B-C) Following initial training on 100% reward, mice were shifted to either 25%, 50%, or 75% reward and trained for 3 days. (B) %Presses met criteria across training. 2-way ANOVA (Probability x Day), no interaction, main effects of Day $F_{3,36} = 6.34$, $p = 0.0015$, and probability group $F_{2,12} = 5.28$, $p = 0.0226$. (C) β coefficients for the interaction between presses that met criteria and n - 1 duration. Interaction between n - 1 duration and n - 1 outcome (Met Yes Reward vs. Met No Reward): $F_{2,17403} = 30.2$; $p < 0.0001$) and 3-way interaction between Duration, Outcome, and Group: $F_{4,17403} = 2.59$; $p = 0.035$. Baseline = Final pretraining day with 100% reward. Prob 1, 2, 3 = Training day 1, 2, or 3 of probability training. RM r = Repeated Measures correlation R. **** $p < 0.0001$, * $p < 0.05$.

**Note S1**, related to Figure 1. As previous reports have indicated that rats in a similarly unstructured task may sometimes make short, stereotyped lever presses after reward (Platt et al., 1973), we investigated our data for evidence of stereotypies. Of the 13 behavioral mice, we found evidence of 1 mouse that appeared to adopt this stereotyped strategy, making presses after a reward that were 450 ± 434ms (mean ± SD) in duration, while the average for all other animals was 1051 ± 757ms. Permutation tests comparing to order shuffled data found that this same mouse exhibited a smaller SD after a rewarded press in actual versus shuffled data on 7 out of 14 days, while no other mouse in this (or subsequent) experiments did so for more than 2 days. Thus, while it is possible for animals to adopt a stereotyped strategy to perform this task, only a very small minority of animals appear to do so. Aside from the difference in species (rats versus mice), the results of Platt et al. (1973) may have been due to the extensive 2 week pretraining period without a duration requirement, wherein rats would have been incentivized to press as rapidly as possible to earn maximal reward and might develop a habitual or stereotyped response that persisted even after introduction of the duration requirement.

**Table S1**. *Simple LME Model Statistics*, related to Figure 2.

| Term | Coef | SE | Upper | Lower | F-Stat | F-Pval | Perm P |
|---|---|---|---|---|---|---|---|
| Intercept | 223 | 49.3 | 320 | 127 | 20.2 | 6.04E-6 | N/A |
| $Dur_{n-1}$ | 0.0909 | 0.00503 | 0.101 | 0.081 | 330 | 1.16E-72 | <0.001 |
| $Dur_{n-2}$ | 0.0731 | 0.00506 | 0.083 | 0.0631 | 207 | 3.64E-47 | <0.001 |
| $Dur_{n-3}$ | 0.0543 | 0.00508 | 0.0642 | 0.0443 | 113 | 1.48E-26 | <0.001 |
| $Dur_{n-4}$ | 0.0376 | 0.0051 | 0.0476 | 0.0276 | 53.9 | 1.88E-13 | <0.001 |
| $Dur_{n-5}$ | 0.0228 | 0.00512 | 0.0328 | 0.0127 | 20.5 | 8.61E-6 | 0.007 |
| $Dur_{n-6}$ | 0.029 | 0.00513 | 0.039 | 0.0189 | 31.1 | 1.63E-8 | <0.001 |
| $Dur_{n-7}$ | 0.0102 | 0.00514 | 0.0203 | 1.78E-4 | 3.62 | 0.0461 | 0.266 |
| $Dur_{n-8}$ | 0.00979 | 0.00515 | 0.0199 | -0.0003 | 3.89 | 0.0572 | 0.307 |
| $Dur_{n-9}$ | 0.0204 | 0.00514 | 0.0305 | 0.0103 | 15.2 | 7.35E-5 | 0.019 |
| $Dur_{n-10}$ | 0.00942 | 0.00513 | 0.0195 | -6.4E-4 | 3 | 0.0664 | 0.328 |
| %Met | 12.6 | 0.533 | 13.7 | 11.6 | 564 | 3.8E-123 | <0.001 |
| Time | 3.44E-5 | 2.84E-6 | 3.99E-5 | 2.88E-5 | 147 | 1.17E-33 | <0.001 |

Parameters, their coefficients, and statistical tests for the simple LME model predicting n duration given n-back durations. Degrees of Freedom for all F-tests = 1, 39711. $Dur_{n-x}$ = Duration of lever press n - x in ms. %Met = %Press Met Criteria. Time = Lever press session timestamp in ms. Coef = β Coefficient. SE = Standard Error. Upper and Lower = 95% confidence intervals. F-stat = F-statistic. F-Pval = p-value from the F-test. Perm P = p-value from permutation test comparing to 1000 order shuffled β coefficients.

**Table S2**. *Complex LME Model Parameters*, related to Figure 2.

| Model Parameters | Description |
|---|---|
| $Dur_{n-1}$ ... $Dur_{n-6}$ | Durations of the prior lever presses from n - 1 to n - 6 |
| MA | A moving average of lever press durations from n - 7 up to n - 60 |
| $HE_{n-1}$ | A binary variable coding for if a checking headentry (HE) was (1) or was not (0) made in between lever press n and press n - 1 |
| $Rew_{n-1}$ | A binary variable coding for if lever press n -1 was (1) or was not (0) rewarded |
| $IPI_{n-1}$ | Inter press interval (IPI, in ms) between lever press n and press n - 1 |
| $IPI_{n-2}$ | As above, only for the IPI between press n and press n - 2 |
| Time | A timestamp (in ms) for when in a session a lever press occurred (from 0ms to 5400000ms) |
| %Met | Overall % of presses that met criteria for a given session |
| $Dur_{n-1}$ : Interactions | Interaction terms between n - 1 Duration and: n - 1 Headentry, n - 1 Reward, n - 1 IPI, Time in Session, and %Met Criteria. Overall question is if the use of recent durations (n - 1) is affected by these variables. |
| MA : Interactions | Interaction terms between the Moving Average (MA) and: n - 1 Headentry, n-1 Reward, n - 1 IPI, Time in Session, and %Met Criteria. Overall question is if the use of the long term moving average is affected by these variables. |

**Table S3**. *Complex LME Model Statistics*, Related to Figure 2.

| Term | Coef | SE | Upper | Lower | F-Stat | Pval |
|---|---|---|---|---|---|---|
| Intercept | 143 | 44 | 229 | 56.3 | 10.5 | 0.0012 |
| *$Dur_{n-1}$* | *-0.0405* | *0.0168* | *-0.00763* | *-0.0735* | *5.83* | *0.0158* |
| $Dur_{n-2}$ | 0.0834 | 0.00609 | 0.0954 | 0.0715 | 188 | 1.29E-42 |
| $Dur_{n-3}$ | 0.0509 | 0.00501 | 0.0608 | 0.0411 | 104 | 2.72E-24 |
| $Dur_{n-4}$ | 0.0351 | 0.00502 | 0.045 | 0.0253 | 48.9 | 2.7E-12 |
| $Dur_{n-5}$ | 0.0188 | 0.00503 | 0.0286 | 0.0089 | 13.9 | 1.91E-4 |
| $Dur_{n-6}$ | 0.0279 | 0.00502 | 0.0377 | 0.0181 | 30.9 | 2.75E-8 |
| MA | 0.354 | 0.0412 | 0.435 | 0.273 | 73.8 | 8.85E-18 |
| $HE_{n-1}$ | -79.5 | 23.1 | -34.3 | -125 | 11.9 | 5.7E-4 |
| $Dur_{n-1}$ : $HE_{n-1}$ | 0.0656 | 0.0118 | 0.0888 | 0.0425 | 30.9 | 2.72E-8 |
| MA : $HE_{n-1}$ | 0.219 | 0.0276 | 0.273 | 0.165 | 62.9 | 2.23E-15 |
| *$Rew_{n-1}$* | *-60.6* | *29.7* | *-2.29* | *-119* | *4.15* | *0.0416* |
| *$Dur_{n-1}$ : $Rew_{n-1}$* | *0.00994* | *0.0151* | *0.0395* | *-0.0196* | *0.436* | *0.509* |
| *MA : $Rew_{n-1}$* | *-0.0414* | *0.0314* | *0.0201* | *-0.103* | *1.74* | *0.187* |
| *$IPI_{n-1}$* | *7.11E-4* | *3.4E-4* | *0.00138* | *4.34E-5* | *4.36* | *0.0368* |
| $Dur_{n-1}$ : $IPI_{n-1}$ | -7.9E-7 | 1.25E-7 | -5.4E-7 | -1E-6 | 39.7 | 2.92E-10 |
| *MA : $IPI_{n-1}$* | *-4.1E-8* | *3.34E-7* | *6.13E-7* | *-7E-7* | *0.0153* | *0.902* |
| *$IPI_{n-2}$* | *2.15E-6* | *1.36E-4* | *2.69E-4* | *-0.00027* | *2.48E-4* | *0.987* |
| $Dur_{n-2}$ : $IPI_{n-2}$ | -3.3E-7 | 7.72E-8 | -1.8E-7 | -4.8E-7 | 18.3 | 1.89E-5 |
| *Time* | 1.51E-5 | 7.58E-6 | 3.0E-5 | 2.85E-7 | 3.99 | 0.0458 |
| $Dur_{n-1}$ : Time | 2.16E-8 | 3.94E-9 | 2.93E-8 | 1.39E-8 | 30 | 4.37E-8 |
| *MA : Time* | *-8.3E-9* | *8.85E-9* | *9.01E-9* | *-2.6E-8* | *0.887* | *0.346* |
| %Met | 10.9 | 1.13 | 13.1 | 8.64 | 92.5 | 7E-22 |
| $Dur_{n-1}$ : %Met | 0.00271 | 4.3E-4 | 0.00355 | 0.00187 | 39.7 | 3E-10 |
| MA : %Met | -0.00491 | 0.00113 | -0.0027 | -0.00713 | 18.9 | 1.42E-5 |

Parameters, their coefficients, and statistical tests for the Complex LME model. Degrees of Freedom for all F-tests = 1, 40203. Some parameters did not improve the model by BIC, but were included in the model because: 1) null effects are of interest (e.g., $Rew_{n-1}$), 2) non-significant main effects were kept if their interactions were significant (e.g., $IPI_{n-2}$) and 3) we kept the same interaction terms for both $Dur_{n-1}$ and the MA to determine if they were differentially controlled by experiential variables (e.g., MA : Time). These BIC non-significant terms are italicized. $Dur_{n-x}$ = Duration of press n - x. MA = Moving Average. $HE_{n-1}$ = Headentry between presses n and n - 1. $Rew_{n-1}$ = n - 1 reward. Data presented here for these two binary variables indicates the effect if they occur (e.g., the interaction between $Dur_{n-1}$ : $HE_{n-1}$ shows how the contribution of Duration changes given that a HE occurred in between press n and press n - 1). $IPI_{n-x}$ = IPI between press n and press n - x. Time = Time in session that a lever press occurred. %Met = % of presses that met the duration criterion for a given session. Coef = β Coefficient. SE = Standard Error. Upper and Lower = 95% confidence intervals. F-stat = F-statistic. Pval = p-value from the F-test.

**Table S4**. *M2 Sham and Lesion Complex LME Model Statistics*, Related to Figure 3.

| | Sham (n = 24630, df = 24605) | | | | Lesion (n = 33348, df = 33323) | | | |
|---|---|---|---|---|---|---|---|---|
| Term | Coef | SE | F | Pval | Coef | SE | F | Pval |
| Intercept | 266 | 58.5 | 20.7 | 5.5E-6 | 186 | 46.3 | 16.1 | 6.08E-5 |
| **$Dur_{n-1}$** | **42.3** | **33** | **1.64** | **0.2** | **0.0708** | **0.0185** | **14.6** | **1.32E-4** |
| $Dur_{n-2}$ | -0.0738 | 0.0221 | 11.1 | 8.59E-4 | 0.0516 | 0.00621 | 69.2 | 9.34E-17 |
| $Dur_{n-3}$ | 0.0523 | 0.00767 | 46.5 | 9.43E-12 | 0.0528 | 0.00552 | 91.4 | 1.24E-21 |
| **$Dur_{n-4}$** | **0.0428** | **0.00638** | **45.1** | **1.88E-11** | **0.0158** | **0.00553** | **8.19** | **0.00422** |
| $Dur_{n-5}$ | 0.0408 | 0.00638 | 40.9 | 1.61E-10 | 0.0302 | 0.00553 | 29.9 | 4.61E-8 |
| $Dur_{n-6}$ | 0.0168 | 0.00639 | 6.94 | 0.00843 | 0.0279 | 0.00551 | 25.6 | 4.14E-7 |
| MA | 0.244 | 0.0566 | 18.6 | 1.66E-5 | 0.216 | 0.0445 | 23.7 | 1.13E-6 |
| $HE_{n-1}$ | -43.4 | 25.3 | 2.93 | 0.0868 | -22.3 | 22.7 | 0.969 | 0.325 |
| **$Dur_{n-1}$ : $HE_{n-1}$** | **0.0635** | **0.0153** | **17.2** | **3.39E-5** | **-0.0149** | **0.0131** | **1.3** | **0.254** |
| MA : $HE_{n-1}$ | 0.149 | 0.032 | 21.8 | 3.01E-6 | 0.165 | 0.0291 | 32.2 | 1.42E-8 |
| $Rew_{n-1}$ | 42.3 | 33 | 1.64 | 0.2 | -41.6 | 31 | 1.8 | 0.179 |
| **$Dur_{n-1}$ : $Rew_{n-1}$** | **-0.065** | **0.0192** | **11.4** | **7.21E-4** | **0.0014** | **0.0168** | **6.91E-3** | **0.934** |
| MA : $Rew_{n-1}$ | -0.101 | 0.0375 | 7.25 | 0.00708 | -0.0354 | 0.0347 | 1.04 | 0.307 |
| $IPI_{n-1}$ | -1.2E-4 | 4.41E-4 | 0.0694 | 0.792 | -1.29E-3 | 4.11E-4 | 9.83 | 0.00172 |
| **$Dur_{n-1}$ : $IPI_{n-1}$** | **-6.7E-07** | **1.6E-7** | **17.7** | **2.58E-5** | **-1.2E-7** | **7.99E-8** | **2.24** | **0.134** |
| MA : $IPI_{n-1}$ | 9E-7 | 4.22E-7 | 4.55 | 0.0329 | 1.63E-6 | 3.98E-7 | 16.7 | 4.29E-5 |
| $IPI_{n-2}$ | -0.0005 | 1.55E-4 | 10.6 | 0.00116 | -4.4E-4 | 1.31E-4 | 11.4 | 7.31E-4 |
| $Dur_{n-2}$ : $IPI_{n-2}$ | -2.3E-7 | 8.2E-8 | 8.05 | 0.00454 | -3.3E-8 | 5.11E-8 | 0.421 | 0.516 |
| Time | 2.33E-5 | 8.27E-6 | 7.93 | 0.00486 | 2.03E-5 | 7.39E-6 | 7.53 | 0.00607 |
| **$Dur_{n-1}$ : Time** | **3.18E-8** | **4.95E-9** | **41.2** | **1.39E-10** | **1.26E-8** | **4.1E-9** | **9.44** | **0.00213** |
| MA : Time | -1.7E-8 | 9.63E-9 | 3.03 | 0.0815 | -1.1E-8 | 8.47E-9 | 1.8 | 0.18 |
| %Met | 8.32 | 1.52 | 30 | 3.97E-8 | 8.03 | 1.18 | 46.6 | 8.82E-12 |
| **$Dur_{n-1}$ : %Met** | **0.00382** | **5.96E-4** | **41** | **1.54E-10** | **8.22E-4** | **4.44E-4** | **3.42** | **0.0644** |
| MA : %Met | -8.6E-4 | 0.0016 | 0.293 | 0.589 | 0.00179 | 0.00119 | 2.28 | 0.131 |

Parameters, their coefficients, and statistical tests for the Complex LME models from M2 Sham and M2 Lesion groups. Bolded terms denote significant Sham/Lesion group differences, assessed using unpaired t-tests with Benjamini-Hockberg false discovery correction as follows. DF for all the following tests are 57976. $Dur_{n-1}$; $t = 5.03$, $p < 0.0001$. $Dur_{n-4}$; $t = 2.96$, $p = 0.003$. $Dur_{n-1}$ : $HE_{n-1}$; $t = 3.89$, $p < 0.0001$. $Dur_{n-1}$ : $Rew_{n-1}$; $t = 2.59$ , $p = 0.00951$. $Dur_{n-1}$ : $IPI_{n-1}$; $t = 3.34$, $p = 0.000845$. $Dur_{n-1}$ : Time; $t = 3.00$, $p = 0.00267$. $Dur_{n-1}$ : %Met; $t = 4.12$ , $p < 0.0001$. $Dur_{n-x}$ = Duration of press n - x. MA = Moving Average. $HE_{n-1}$ = Headentry between presses n and n - 1. $Rew_{n-1}$ = n - 1 reward. $IPI_{n-x}$ = IPI between press n and press n - x. Time = Time in session that a lever press occurred. %Met = % of presses that met the duration criterion for a given session. Coef = β Coefficient. SE = Standard Error. Upper and Lower = 95% confidence intervals. F = F-statistic. Pval = p-value from the F-test.

**Table S5**, *M2 GCaMP Complex LME Model Statistics*, related to Figure 4.

| | Before Press, df = 11359 | | | During Press, df = 6759 | | | After Press, df = 11359 | | |
|---|---|---|---|---|---|---|---|---|---|
| Term | Coef | F | p | Coef | F | p | Coef | F | p |
| Int. | 2.26 | 1.47 | 0.225 | 0.82 | 0.131 | 0.718 | -1.76 | 0.704 | 0.402 |
| Dur$_{n}$ | **-4.0E-4** | **4.86** | **0.0275** | **-1.95E-3** | **75.6** | **4.26E-18** | -2.3E-4 | 0.249 | 0.618 |
| Dur$_{n-1}$ | **-1.6E-3** | **5.34** | **0.0209** | 9.04E-4 | 0.933 | 0.334 | -3.2E-4 | 0.198 | 0.656 |
| Dur$_{n-2}$ | **6.35E-4** | **9.16** | **0.00248** | **6.21E-4** | **5.6** | **0.018** | **5.24E-4** | **5.75** | **0.0165** |
| Dur$_{n-3}$ | 1.0E-4 | 0.302 | 0.583 | -2.6E-4 | 1.35 | 0.245 | 2.3E-4 | 1.46 | 0.226 |
| Dur$_{n-4}$ | 2.61E-4 | 2.02 | 0.155 | **4.62E-4** | **4.15** | **0.0416** | 2.77E-4 | 2.1 | 0.147 |
| Dur$_{n-5}$ | 6.81E-5 | 0.137 | 0.711 | -4.7E-5 | 0.0437 | 0.834 | 7.87E-5 | 0.169 | 0.681 |
| Dur$_{n-6}$ | 2.1E-4 | 1.31 | 0.253 | 2.5E-4 | 1.22 | 0.269 | 3.21E-4 | 2.8 | 0.0942 |
| MA | -1.49E-3 | 0.699 | 0.403 | 2.2E-4 | 0.0086 | 0.926 | 0.00138 | 0.547 | 0.46 |
| HE$_{n-1}$ | **8.02** | **25.2** | **5.3E-7** | **8.29** | **14.8** | **1.19E-4** | 2.45 | 2.15 | 0.142 |
| Dur$_{n-1}$ : HE$_{n-1}$ | **0.00241** | **24.6** | **7.13E-7** | **0.00217** | **13.4** | **2.48E-4** | **0.00269** | **28.1** | **1.15E-7** |
| MA : HE$_{n-1}$ | **-6.27E-3** | **14.6** | **1.35E-4** | **-5.91E-3** | **8.15** | **0.00433** | -2.88E-3 | 2.84 | 0.0919 |
| Rew$_{n-1}$ | -1.65 | 0.686 | 0.407 | -0.0339 | 1.79E-4 | 0.989 | 1.03 | 0.247 | 0.619 |
| Dur$_{n-1}$ : Rew$_{n-1}$ | **0.00241** | **14.6** | **1.34E-4** | 0.00128 | 2.49 | 0.115 | 1.22E-4 | 0.0346 | 0.852 |
| MA : Rew$_{n-1}$ | 1.13E-4 | 0.00482 | 0.945 | 01.98E-4 | 0.00987 | 0.921 | 0.00197 | 1.37 | 0.242 |
| IPI$_{n-1}$ | **7.05E-5** | **15.8** | **7.14E-5** | **5.86E-5** | **6.59** | **0.0103** | **6.39E-5** | **12** | **5.4E-4** |
| Dur$_{n-1}$ : IPI$_{n-1}$ | **-1.3E-8** | **6.07** | **0.0138** | -1.2E-8 | 3.72 | 0.0537 | **-1.5E-8** | **7.21** | **0.00727** |
| MA : IPI$_{n-1}$ | -2.8E-8 | 3.34 | 0.0678 | -2.2E-8 | 1.26 | 0.263 | -2.3E-8 | 2.04 | 0.154 |
| IPI$_{n-2}$ | 2.19E-6 | 0.0729 | 0.787 | 7.99E-6 | 0.702 | 0.402 | 5.4E-6 | 0.41 | 0.522 |
| Dur$_{n-2}$ : IPI$_{n-2}$ | -4.9E-9 | 1.04 | 0.308 | **-1.2E-8** | **4.64** | **0.0313** | **-1E-8** | **4.24** | **0.0395** |
| Time | **-1.2E-6** | **9.62** | **0.00193** | -7.1E-7 | 1.69 | 0.194 | **-1.2E-6** | **9.96** | **0.0016** |
| Dur$_{n-1}$ : Time | -7E-11 | 0.234 | 0.628 | -1.7E-10 | 0.843 | 0.359 | -1E-10 | 0.468 | 0.494 |
| MA : Time | **1.01E-9** | **5.48** | **0.0192** | 7.68E-10 | 1.7 | 0.193 | **1.3E-9** | **8.37** | **0.00382** |
| %Met | -0.025 | 0.174 | 0.676 | -0.0268 | 0.107 | 0.743 | 0.0409 | 0.412 | 0.521 |

| | | | | | | | | | |
|---|---|---|---|---|---|---|---|---|---|
| $Dur_{n-1}$ : %Met | -3.2E-5 | 1.77 | 0.184 | -5.6E-5 | 3.48 | 0.0621 | -1.4E-5 | 0.293 | 0.588 |
| MA : %Met | 5.27E-5 | 0.913 | 0.339 | 6.09E-5 | 0.727 | 0.394 | -4.2E-5 | 0.491 | 0.483 |
| $Act_{n-1}$ | **0.453** | **2370** | **0** | **0.393** | **1070** | **1.3E-217** | **0.431** | **2130** | **0** |
| $Act_{n-2}$ | -3.77E-3 | 0.138 | 0.711 | 0.0202 | 2.48 | 0.115 | 0.0106 | 1.1 | 0.295 |
| $Act_{n-3}$ | **0.0251** | **6.07** | **0.0138** | -0.0186 | 2.09 | 0.148 | -3.66E-3 | 0.131 | 0.717 |
| $Act_{n-4}$ | -5.33E-3 | 0.274 | 0.601 | 0.0208 | 2.59 | 0.108 | **0.0481** | **22.6** | **2.0E-6** |
| $Act_{n-5}$ | 0.0134 | 1.74 | 0.187 | 0.00597 | 0.216 | 0.642 | **-0.0215** | **4.52** | **0.0336** |
| $Act_{n-6}$ | 0.0129 | 1.93 | 0.165 | 0.017 | 2.01 | 0.157 | **0.0259** | **7.8** | **0.00524** |
| $Rew_n$ | NA | NA | NA | NA | NA | NA | **5.79** | **13.4** | **2.52E-4** |
| $Dur_n$ : $Rew_n$ | NA | NA | NA | NA | NA | NA | 8.35E-4 | 2.04 | 0.153 |
| MA : $Rew_n$ | NA | NA | NA | NA | NA | NA | **-7.64E-3** | **29.8** | **4.99E-8** |

Parameters, their coefficients, and statistical tests relating M2 calcium activity to behavior. We predicted activity at three different time points -1s to 0s Before Press, During the Press, and 0s to +1s After Press offset. We included prior activity ($Act_{n-x}$) as a covariate to control for autocorrelation in calcium activity data. Bolded terms are significant by F-test on the model. In the After Press model, we also incorporated an n - 0 reward term (i.e., was the just completed press rewarded) given that we see an apparent reward response at this time point (Figure 4D). P-values of 0 are reported for some prior activity terms due to Matlab's numerical resolution. $Dur_{n-x}$ = Duration of press n - x. MA = Moving Average. $HE_{n-1}$ = Headentry between presses n and n - 1. $Rew_{n-1}$ = n - 1 reward. $IPI_{n-x}$ = IPI between press n and press n - x. Time = Time in session that a lever press occurred. %Met = % of presses that met the duration criterion for a given session. Coef = β Coefficient. SE = Standard Error. Upper and Lower = 95% confidence intervals. F = F-statistic. p = p-value from the F-test.

**Table S6**. *M2-DMS GCaMP Complex LME Model Statistics*, related to Figure 5.

| | Before Press, df = 12038 | | | During Press, df = 7524 | | | After Press, df = 12025 | | |
|---|---|---|---|---|---|---|---|---|---|
| Term | Coef | F | p | Coef | F | p | Coef | F | p |
| Int. | -0.501 | 0.0655 | 0.798 | -0.461 | 0.0273 | 0.869 | -2.06 | 0.815 | 0.367 |
| Dur$_{n-0}$ | **6.79E-4** | **11.2** | **8.22E-4** | -3.1E-4 | 1.42 | 0.234 | **0.00247** | **16.8** | **4.15E-5** |
| Dur$_{n-1}$ | -1.48E-3 | 3.22 | 0.0727 | 8.22E-5 | 0.00508 | 0.943 | -1.49E-3 | 2.58 | 0.108 |
| Dur$_{n-2}$ | -2.4E-5 | 0.0102 | 0.92 | 1.69E-4 | 0.283 | 0.594 | 4.82E-5 | 0.0318 | 0.858 |
| Dur$_{n-3}$ | 6.01E-6 | 8.58E-4 | 0.977 | -3.1E-5 | 0.0134 | 0.908 | 1.36E-4 | 0.349 | 0.554 |
| Dur$_{n-4}$ | -1.4E-4 | 0.435 | 0.51 | -6.1E-5 | 0.0526 | 0.819 | 3.41E-4 | 2.19 | 0.139 |
| Dur$_{n-5}$ | -1.1E-4 | 0.261 | 0.61 | -1.6E-4 | 0.348 | 0.555 | -9.6E-6 | 0.00172 | 0.967 |
| Dur$_{n-6}$ | -4.3E-5 | 0.0426 | 0.837 | 2.37E-4 | 0.789 | 0.375 | -9.8E-5 | 0.179 | 0.672 |
| MA | -2.12E-3 | 1.08 | 0.298 | -5.08E-3 | 3.06 | 0.0804 | -3.5E-3 | 2.33 | 0.127 |
| HE$_{n-1}$ | **4.49** | **4.29** | **0.0382** | **6.41** | **4.09** | **0.0432** | 0.751 | 0.0957 | 0.757 |
| Dur$_{n-1}$ : HE$_{n-1}$ | **0.00126** | **4.91** | **0.0268** | **0.00213** | **8.67** | **0.00325** | **0.00136** | **4.53** | **0.0334** |
| MA : HE$_{n-1}$ | 5.42E-5 | 6.85E-4 | 0.979 | -2.34E-3 | 0.627 | 0.429 | 0.00404 | 3.02 | 0.0821 |
| Rew$_{n-1}$ | **6.91** | **7.19** | **0.00732** | 5.72 | 2.69 | 0.101 | **9.31** | **10.4** | **0.00127** |
| Dur$_{n-1}$ : Rew$_{n-1}$ | -9.5E-4 | 1.67 | 0.196 | -6.3E-4 | 0.425 | 0.514 | 2.71E-4 | 0.108 | 0.742 |
| MA : Rew$_{n-1}$ | -1.55E-3 | 0.514 | 0.473 | -5.6E-4 | 0.0371 | 0.847 | -2.79E-3 | 1.34 | 0.248 |
| IPI$_{n-1}$ | **-7.2E-5** | **13.6** | **2.3E-4** | **-6.7E-5** | **4.76** | **0.0292** | **-8.8E-5** | **16.2** | **5.66E-5** |
| Dur$_{n-1}$ : IPI$_{n-1}$ | -1.7E-9 | 0.0838 | 0.772 | -3.3E-9 | 0.178 | 0.673 | 7.63E-10 | 0.0136 | 0.907 |
| MA : IPI$_{n-1}$ | **1.17E-7** | **40.3** | **2.25E-10** | **1.21E-7** | **17.4** | **3.12E-5** | **1.01E-7** | **23.9** | **1.0E-6** |
| IPI$_{n-2}$ | -1.5E-5 | 3.21 | 0.0732 | 6.6E-7 | 0.00335 | 0.954 | -4.5E-6 | 0.238 | 0.626 |
| Dur$_{n-2}$ : IPI$_{n-2}$ | 2.62E-9 | 0.32 | 0.572 | 4.26E-9 | 0.462 | 0.497 | 8.68E-11 | 2.8E-4 | 0.987 |
| Time | **1.31E-6** | **8.42** | **0.00371** | -2E-8 | 8.36E-4 | 0.977 | **1.37E-6** | **7.28** | **0.00698** |
| Dur$_{n-1}$ : Time | -1.3E-10 | 0.695 | 0.405 | -9.8E-11 | 0.238 | 0.626 | -3.3E-10 | 3.74 | 0.0531 |
| MA : Time | **-1.5E-9** | **8.25** | **0.00409** | 1.03E-11 | 1.96E-4 | 0.989 | **-1.3E-9** | **5.45** | **0.0196** |
| %Met | 0.0512 | 0.282 | 0.595 | 0.194 | 2.22 | 0.136 | -0.0109 | 0.00966 | 0.922 |

| | | | | | | | | | |
|---|---|---|---|---|---|---|---|---|---|
| Dur$_{n-1}$ : %Met | 5.03E-5 | 3.06 | 0.0803 | -3.3E-5 | 0.784 | 0.376 | 2.22E-6 | 0.00471 | 0.945 |
| MA : %Met | 7.62E-5 | 0.884 | 0.347 | 2.64E-5 | 0.056 | 0.813 | **1.93E-4** | **4.28** | **0.0387** |
| Act$_{n-1}$ | **0.387** | **1850** | **0** | **0.342** | **893** | **1.4E-185** | **0.321** | **1250** | **1.8E-260** |
| Act$_{n-2}$ | **0.0254** | **7.02** | **0.00806** | **0.0322** | **7.25** | **0.00712** | **0.0298** | **9.89** | **0.00167** |
| Act$_{n-3}$ | **0.0316** | **10.9** | **9.72E-4** | **0.0431** | **13** | **3.2E-4** | 0.0162 | 2.94 | 0.0866 |
| Act$_{n-4}$ | **0.0438** | **21** | **4.56E-6** | **0.0384** | **10.3** | **0.00131** | **0.0213** | **5.08** | **0.0242** |
| Act$_{n-5}$ | 0.0111 | 1.34 | 0.247 | 0.0202 | 2.85 | 0.0913 | -9.24E-3 | 0.96 | 0.327 |
| Act$_{n-6}$ | **0.0465** | **27.5** | **1.64E-7** | **0.0653** | **33.5** | **7.59E-9** | **0.0418** | **21.7** | **3.24E-6** |
| Rew$_n$ | NA | NA | NA | NA | NA | NA | **10.1** | **19** | **1.31E-5** |
| Dur$_n$ : Rew$_n$ | NA | NA | NA | NA | NA | NA | **-3.22E-3** | **18.1** | **2.06E-5** |
| MA : Rew$_n$ | NA | NA | NA | NA | NA | NA | **-7.16E-3** | **12.4** | **4.29E-4** |

Parameters, their coefficients, and statistical tests relating M2 calcium activity to behavior. We predicted activity at three different time points -1s to 0s Before Press, During the Press, and 0s to +1s After Press offset. We included prior activity (Act$_{n-x}$) as a covariate to control for autocorrelation in calcium activity data. Bolded terms are significant by F-test on the model. In the After Press model, we also incorporated an n - 0 reward (i.e., was the just completed press rewarded) term given that we see an apparent reward response at this time point (Figure 5D). P-values of 0 are reported for prior activity due to Matlab's numerical resolution. Dur$_{n-x}$ = Duration of press n - x. MA = Moving Average. HE$_{n-1}$ = Headentry between presses n and n - 1. Rew$_{n-1}$ = n - 1 reward. IPI$_{n-x}$ = IPI between press n and press n - x. Time = Time in session that a lever press occurred. %Met = % of presses that met the duration criterion for a given session. Coef = β Coefficient. SE = Standard Error. Upper and Lower = 95% confidence intervals. F = F-statistic. p = p-value from the F-test.

**Table S7**. *M2-DMS Sham and Lesion Complex LME Model Statistics*, related to Figure 6.

| | Sham (n = 23758 df = 23733) | | | | Lesion (n = 23596, df = 23571) | | | |
|---|---|---|---|---|---|---|---|---|
| Term | Coef | SE | F | Pval | Coef | SE | F | Pval |
| Int. | 291 | 55 | 28 | 1.2E-7 | 285 | 58 | 24.1 | 9.17E-7 |
| $Dur_{n-1}$ | -0.0284 | 0.0212 | 1.8 | 0.18 | -0.0313 | 0.0217 | 2.08 | 0.149 |
| $Dur_{n-2}$ | 0.0607 | 0.00763 | 63.3 | 1.83E-15 | 0.0691 | 0.00772 | 80.2 | 3.68E-19 |
| $Dur_{n-3}$ | 0.0229 | 0.00655 | 12.3 | 4.64E-4 | 0.0434 | 0.0066 | 43.2 | 5.03E-11 |
| $Dur_{n-4}$ | 0.0199 | 0.00656 | 9.25 | 0.00235 | 0.0308 | 0.00661 | 21.7 | 3.13E-6 |
| $Dur_{n-5}$ | 0.0301 | 0.00657 | 21 | 4.69E-7 | 0.0415 | 0.00662 | 39.3 | 3.79E-10 |
| $Dur_{n-6}$ | 0.0282 | 0.00658 | 18.3 | 1.86E-6 | 0.017 | 0.00664 | 6.54 | 0.0105 |
| MA | 0.217 | 0.0561 | 15 | 1.08E-4 | 0.2 | 0.0585 | 11.7 | 6.23E-4 |
| $HE_{n-1}$ | -79.1 | 28.6 | 7.66 | 0.00565 | -38.2 | 31.2 | 1.5 | 0.221 |
| **$Dur_{n-1}$ : $HE_{n-1}$** | **0.0295** | **0.0157** | **3.52** | **0.0607** | **-0.0412** | **0.0165** | **6.22** | **0.0126** |
| MA : $HE_{n-1}$ | 0.0959 | 0.0366 | 6.86 | 0.00882 | 0.157 | 0.0399 | 15.6 | 7.94E-5 |
| $Rew_{n-1}$ | -73.2 | 37.7 | 3.77 | 0.0523 | -77.3 | 37.5 | 4.25 | 0.0392 |
| $Dur_{n-1}$ : $Rew_{n-1}$ | -0.0349 | 0.0194 | 3.22 | 0.0725 | -0.00192 | 0.0193 | 0.00983 | 0.921 |
| MA : $Rew_{n-1}$ | 0.0708 | 0.0432 | 2.68 | 0.101 | 0.112 | 0.0464 | 5.8 | 0.0161 |
| $IPI_{n-1}$ | 0.00187 | 4.68E-4 | 16 | 6.32E-5 | 7.19E-4 | 4.03E-4 | 3.18 | 0.0748 |
| $Dur_{n-1}$ : $IPI_{n-1}$ | -6.3E-7 | 1.49E-7 | 17.9 | 2.36E-5 | -5.2E-7 | 1.62E-7 | 10.4 | 0.00129 |
| MA : $IPI_{n-1}$ | -8.3E-7 | 4.75E-7 | 3.04 | 0.0813 | -2.4E-8 | 4.04E-7 | 0.00362 | 0.952 |
| $IPI_{n-2}$ | -4.8E-4 | 1.63E-4 | 8.79 | 0.00304 | -4.3E-4 | 1.59E-4 | 7.25 | 0.00709 |
| $Dur_{n-2}$ : $IPI_{n-2}$ | -5.8E-8 | 9.35E-8 | 0.391 | 0.532 | -1.2E-7 | 9.64E-8 | 1.46 | 0.227 |
| Time | -5.2E-6 | 1.02E-5 | 0.261 | 0.61 | -9.2E-7 | 9.78E-6 | 0.00877 | 0.925 |
| $Dur_{n-1}$ : Time | 3.59E-8 | 4.93E-9 | 52.9 | 3.59E-13 | 3.39E-8 | 5.04E-9 | 45.2 | 1.81E-11 |
| MA : Time | 8.68E-9 | 1.17E-8 | 0.553 | 0.457 | 1.43E-9 | 1.21E-8 | 0.014 | 0.906 |
| %Met | 7.95 | 1.37 | 33.8 | 6.29E-9 | 7.74 | 1.55 | 25 | 5.7E-7 |
| $Dur_{n-1}$ : %Met | 0.00276 | 5.12E-4 | 29 | 7.22E-8 | 0.00105 | 6.01E-4 | 3.06 | 0.0804 |
| MA : %Met | -0.0013 | 0.00146 | 0.792 | 0.374 | -1.3E-4 | 0.00172 | 0.00543 | 0.941 |

Parameters, their coefficients, and statistical tests for the Complex LME models from M2-DMS Sham and Lesion groups. Bolded terms denote significant Sham/Lesion group differences, assessed using unpaired t-tests with Benjamini-Hockberg false discovery correction. Only the $Dur_{n-1}$ : $HE_{n-1}$ interaction significantly differed between Sham and Lesion; $t_{47352} = 3.10$, $p = 0.00193$. $Dur_{n-x}$ = Duration of press n - x. MA = Moving Average. $HE_{n-1}$ = Headentry between presses n and n - 1. $Rew_{n-1}$ = n - 1 reward. $IPI_{n-x}$ = IPI between press n and press n - x. Time =

Time in session that a lever press occurred. %Met = % of presses that met the duration criterion for a given session. Coef = β Coefficient. SE = Standard Error. Upper and Lower = 95% confidence intervals. F = F-statistic. Pval = p-value from the F-test.
